# Tyrosine phosphorylation coupling of one carbon metabolism and virulence in an endogenous pathogen

**DOI:** 10.1101/2025.03.11.642667

**Authors:** Kendall S. Stocke, Satya D. Pandey, Shunying Jin, John D. Perpich, Lan Yakoumatos, Hirotaka Kosaki, Daniel W. Wilkey, Zackary R. Fitzsimonds, Aruna Vashishta, Ian Snider, Mukesh K. Sriwastva, Hong Li, Jiu-Zhen Jin, Daniel P. Miller, Michael L. Merchant, Juhi Bagaitkar, Silvia M. Uriarte, Jan Potempa, Richard J. Lamont

## Abstract

Endogenous pathogens can constrain virulence to ensure survival in the host. Pathogenic state can be controlled by metabolic responses to the prevailing microenvironment; however, the coupling and effector mechanisms are not well understood. Flux through the One Carbon Metabolism (OCM) pathway can modulate virulence of the oral pathobiont *Porphyromonas gingivalis*, and here we show that this is controlled by tyrosine phosphorylation-dependent differential partitioning of gingipain proteases. The OCM essential precursor pABA inhibits the low molecular weight tyrosine phosphatase Ltp1, and consequently relieves inhibition of its cognate kinase, Ptk1. We found that in the absence of pABA, reduced Ptk1 kinase activity blocks extracellular release of gingipains. Surface retention of gingipains confers resistance to neutrophil mobilization and killing, and virulence in animal models of disease is elevated. Reciprocally, Ptk1 and gingipains are required for maximal flux through OCM, and Ptk1 can phosphorylate the OCM pathway enzymes GlyA and GcvT. Further, ALP, an alkaline phosphatase involved in synthesis of DHPPP, which combines with pABA to make DHP, is phosphorylated and activated by Ptk1. We propose, therefore, that although the primary function of Ptk1 is to maintain OCM balance, it mechanistically couples metabolism with tunable pathogenic potential through directing the location of proteolytic virulence factors.

## Introduction

The human microbiome, while normally symbiotic, contains organisms with context-dependent pathogenic potential; pathobionts that can transition to a virulent state in response to changing environmental influences. In many cases, the virulence factors of these pathobionts are well-characterized. Less understood, however, are the mechanisms which suppress virulence the majority of the time, and function to maintain a commensal relationship with the host. What has become evident is that pathogenicity is one outcome of the integration of environmental cues with bacterial metabolic programming to optimize survival and nutrient availability (1–3). One Carbon Metabolism (OCM) is an essential component of cellular intermediary metabolism, producing a number of one-carbon unit intermediates (formyl, methylene, methenyl, methyl) which are required for the synthesis of various amino acids and other biomolecules such as purines, thymidylate, and redox regulators (4, 5). OCM can play an over-riding role in microbial pathogenicity in that auxotrophy in para-aminobenzoic acid (pABA), an essential OCM precursor, results in an inability of some microorganisms to survive in vivo due to the low levels of pABA present in mammalian systems (6, 7). However, the interrelationships between pABA and virulence can be more nuanced, for example in *Vibrio fischeri* pABA induces an increase in c-di-GMP, which is necessary to induce biofilm formation (8). We have found that the asaccharolytic periodontal pathobiont *Porphyromonas gingivalis* can scavenge pABA from partner species such as *Streptococcus gordonii*, and in the absence of exogenous pABA, the virulence of *P. gingivalis* increases (9). As *P. gingivalis* is non-pathogenic in murine models without support from a polymicrobial community (10), OCM and pABA are positioned as regulators of context-dependent virulence in the organism. However, the pathogenic program that responds to pABA availability and the coupling between OCM and virulence are largely unexplored. Here we uncover a tyrosine phosphorylation-based molecular couple which links OCM with secretion of gingipain protease virulence factors in *P. gingivalis*.

## Results

### PabC regulates virulence and tyrosine phosphorylation

To isolate and define the role of pABA in virulence we deleted *pabC* encoding the PabC enzyme, a 4-amino-4-deoxychorismate lyase which synthesizes pABA from 4-amino 4 deoxychorismate (11). We then utilized a germ-free mouse model to test the pathogenicity of *P. gingivalis* in the absence of inter-species interactions, and avoiding chemical complementation by pABA produced by the endogenous microbiota. In this model, the *P. gingivalis* parental strain exhibited no significant alveolar bone loss (Fig. 1a) consistent with the previously reported community-dependent pathogenicity of the organism (10). In contrast, monoinfection with the *ΔpabC* mutant induced periodontal alveolar bone loss (Fig. 1a), indicating that pABA deficiency mimics the community-dependent pathogenic state of the organism. A comparison of the expression of classical virulence-associated genes, including those encoding gingipains, fimbriae and hemin uptake systems, did not reveal differential expression in the Δ*pabC* mutant (Fig. 1b), suggestive of a novel relationship between OCM and virulence in *P. gingivalis*. As an amino-benzoate, pABA is a competitive inhibitor of low molecular weight protein tyrosine phosphatases (LMW-PTPs) (12), including Ltp1 in *P. gingivalis* (13). Ltp1 is a cognate phosphatase and inhibitor of the multifunctional bacterial tyrosine (BY) kinase Ptk1 (14), and thus we turned our attention to the phosphorylation status of Ptk1 in Δ*pabC*. FLAG-Ptk1 was expressed in Δ*pabC* and WT strains, and following precipitation with FLAG antibodies, blots were probed with phosphotyrosine antibodies. In the absence of PabC, the phosphorylation status of Ptk1 was reduced, and this inhibition of Ptk1 phosphorylation was reversed by the addition of exogenous pABA (Fig. 1c). We postulated, therefore, that a major mode of action of pABA in relation to virulence is in the regulation of Ptk1 activity, and hence loss of Ptk1 would be predicted to phenocopy the Δ*pabC* mutant and cause an increase in virulence. This was confirmed in the alveolar bone loss model, where Δ*ptk1* was more pathogenic than the WT strain in conventional animals (as the absence of Ptk1 obviates the need to restrict pABA availability) (Fig. 1d).

**Figure 1.**
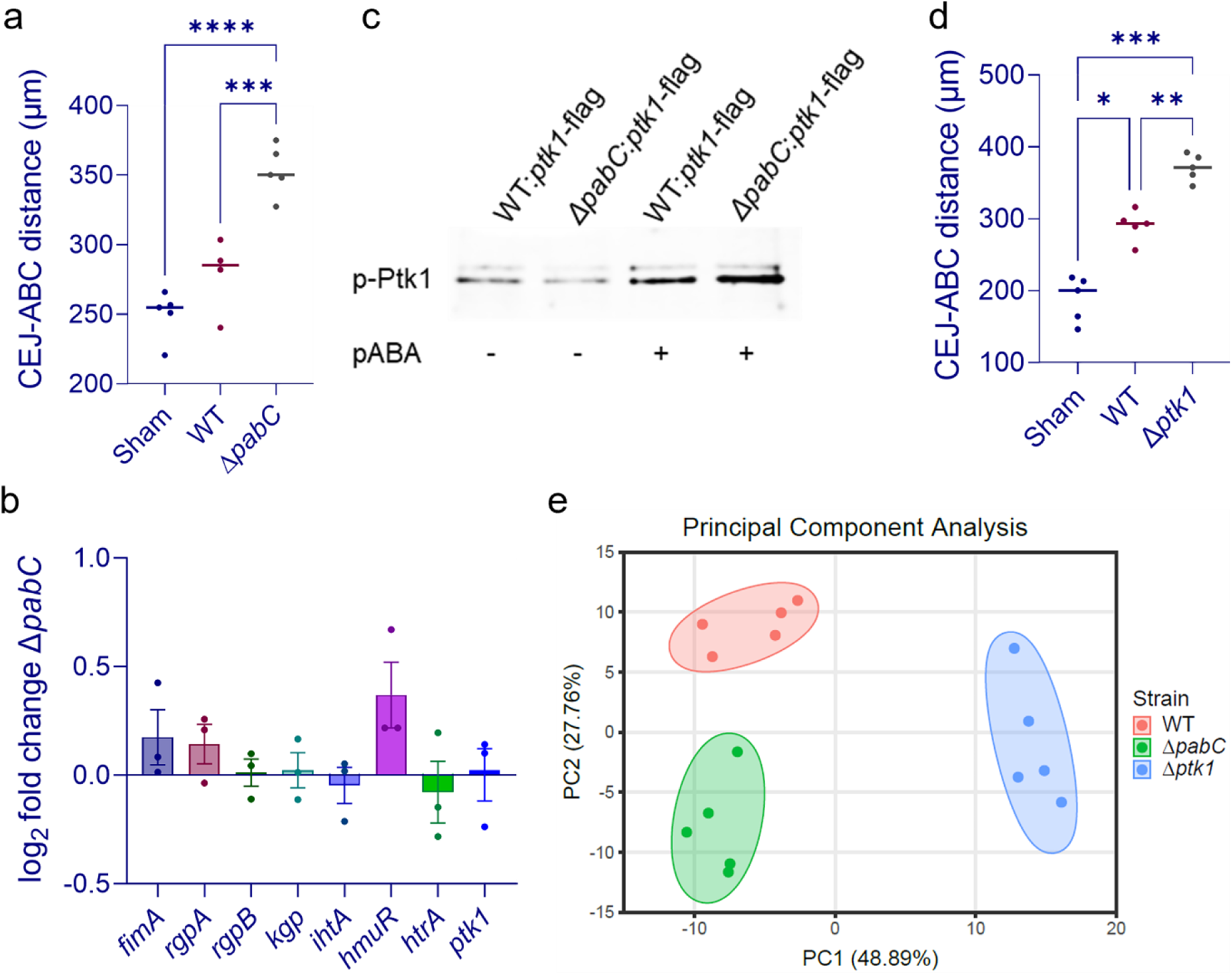
pABA regulated the phosphorylation status of Ptkk1 and mutants lacking PabC or Ptk1 are more pathogenic than WT. a) and d) μCT analysis of alveolar bone loss in germ-free (a) or conventional (d) mice following infection with the *P. gingivalis* strains indicated, or sham infected. Reconstructed images of the maxillary molars were analyzed along the sagittal slice to determine the distance between the ABC (alveolar bone crest) and the CEJ (cementoenamel junction). The data are the means of 6 interdental points at 3 sites around the first and second maxillary molars in each animal. \**P*<.05, \*\**P*<.01, \*\*\**P*<.005, \*\*\*\**P*<.001 relative to sham infection by one-way analysis of variance (ANOVA) with Tukey’s correction for multiple comparisons. b) qRT-PCR of mRNA levels of genes indicated by the 2^−ΔΔCt^ method and with 16s rRNA as the internal control. Data are expressed as Log_2_ fold change in Δ*pabC* relative to WT and are representative of three biological replicates. There was no significant difference (*P*>.05 ANOVA with Tukey’s correction for multiple comparisons) between Δ*pabC* and WT for any of the genes tested. c) Western blot of FLAG antibody immunoprecipitate of lysates of strains indicated. Blots were probed with phosphotyrosine antibodies. Strains were incubated with pABA 0.25 mg/ml where indicated. Image is representative of three biological replicates. e) Principal Component Analysis (PCA) of RNASeq data from WT and Δ*pabC* and Δ*ptk1* strains.

### Transcriptional regulation through PabC and Ptk1

To further investigate the pathogenic characteristics of the mutant strains, we utilized a murine subcutaneous infection model and performed an RNA-Seq analysis on organisms recovered from abscesses. Principal Component Analysis (Fig.1e) of the transcriptome showed that the two mutant strains clustered separately, and each was distinct from the WT parental strain. Distinct transcriptional profiles of WT and mutant strains were also apparent in heat map and Venn diagram of differentially expressed genes (DEGs) (Supplementary Data S1, Fig. S1 a,b). Examination of DEGs at a Log2 fold change of 1 and an adjusted *P* < .05 (Fig. S1 c,d, Supplementary Data S2) did not reveal transcriptional patterns consistent with regulation of virulence. Consideration of all statistically significantly down-regulated genes (598) without regard to fold change (Supplementary Data S1), revealed that 16 were related to OCM (Fig. S2) consistent with a reduction in flux through OCM in the absence of the precursor pABA. Nevertheless, these low levels of differential expression (Log_2_ 0.6-0.7) also suggest that transcriptional control is not the primary means by which OCM flux is regulated. Of note, *ltp1* was down-regulated in Δ*pabC*, potentially a transcriptional response to minimize the effect of loss of an Ltp1 inhibitor. In Δ*ptk1* there was downregulation of the *fimA-E* operon, encoding the major fimbrial adhesin structure and a documented component of the tyrosine phosphorylation signaling network (15). With both mutants, a high percentage of regulated genes was unannotated. Operon prediction using *Rockhopper* (16) identified an operon spanning PGN_1067-1074, along with the immediately upstream gene (PGN_1075) transcribed in the opposite direction, that were upregulated in Δ*ptk1* (although the Log2 differential expression of PGN_1074, while statistically significant, was 0.9, just below our threshold of 1.0, Supplementary Data S1). In Δ*pabC*, PGN_1067-1072 and PGN_1075 were also upregulated, while PGN_1073-1074 significantly trended upward (Supplementary Data S1 and S2). PGN_1069 has domain homologies with aspartate, glutamate and uridylate kinase families, and PGN_1071 has a predicted tyrosine kinase domain, allowing speculation that upregulation of this operon may be a feedback-mediated adjustment to the reduction of Ptk1 BY kinase activity. Moreover, we have previously reported that PGN_1071 and PGN_1073 are conditionally essential for *P. gingivalis* fitness in a murine abscess model and for a sustained interaction with gingival epithelial cells (17). Collectively, these data suggest that indirect activation of Ptk1 by pABA may be partial, and while sufficient to regulate virulence, does not impact all other pathways controlled by tyrosine phosphorylation/dephosphorylation at the transcriptional level, especially given the presence of other kinases and phosphatases in *P. gingivalis* (18). Similarly, disruption of OCM can have effects independent of Ptk1. STRING functional enrichment analysis showed the major transcriptional impact of loss of PabC, among GO Biological Processes, was upregulation of CRISPR-associated antiviral responses (Fig. S1e). CRISPR systems are recognized as involved in *P. gingivalis* virulence through regulation of a putative auto-transporter adhesin (19). Although the corresponding gene for this adhesin, PGN_1547, was not identified as differentially regulated, CRISPR systems may control expression of other novel virulence factors and contribute to PabC-associated virulence. GO Biological Processes analysis of differentially regulated genes in Δ*ptk1*, which can no longer produce extracellular polysaccharide (EPS) (14), showed downregulation of genes involved in this biosynthetic pathway (Fig. S1f).

### Posttranscriptional control of virulence factors by the PabC-Ptk1 axis

The absence of a clear connection between the transcriptional landscape and virulence of the *pabC* and *ptk1* mutants led us to consider a posttranslational axis. We recently reported that optimal processing of the gingipain proteases RgpA, RgpB and Kgp, major virulence factors of *P. gingivalis*, requires tyrosine phosphorylation (20). We used blotting of cell fractions to investigate localization of the gingipains in Δ*pabC* and Δ*ptk1*. Coomassie stained gels of the fractions are shown in Fig. S3a, b. In both mutants, there was an increase in retention of RgpA/B and Kgp in the membrane fractions, although posttranslational processing was not affected (Fig. 2a). proRgpB was processed to the membrane type gingipain (mt-RgpB, ∼75 kDa) consisting of the catalytic domain (∼55 kDa) and the Ig-like domain to which A-LPS attaches on the C-terminal residue. Similarly, proRgpA and proKgp were correctly processed into the non-covalent complex of catalytic and hemagglutinin/adhesion (H/A) anchored into the outer membrane with A-LPS attached to C-terminal H/A domain (21). Correspondingly, in the Δ*pabC* and Δ*ptk1* mutants, release of soluble proteolytically processed, A-LPS-free, gingipains into the medium was reduced. Absence of Ptk1 or PabC does not cause a general secretion defect in *P. gingivalis*, as, excluding gingipains, the overall profile of proteins secreted into the extracellular milieu was similar to WT (Fig. S3c). These results suggest that in the absence of tyrosine phosphorylation, active gingipains remain associated with A-LPS in the membranes. In support of this concept, we found increased levels of cell surface A-LPS in Δ*pabC* and Δ*ptk1* compared to the parental strain (Fig. 2b, c). A-LPS is conjugated to gingipains by the PorU sortase (22), and the *porU* gene was down-regulated in Δ*pabC* (Fig. S1c, Supplementary Data S2). Similar to the reduction in *ltp1* expression, this may be a transcriptional response to compensate for increased A-LPS on the cell surface which is insufficient to significantly impact phenotypic properties. Further, enzyme assays confirmed increased cell-associated gingipain activity in both mutant strains with reduced activity in the culture supernatants (Fig. 2d-g). Genetic complementation with the relevant wild type allele in trans and, in the case of Δ*pabC*, chemical complementation with exogenous pABA, reversed the effect of the gene deletion (Fig. 2h-k).

**Figure 2.**
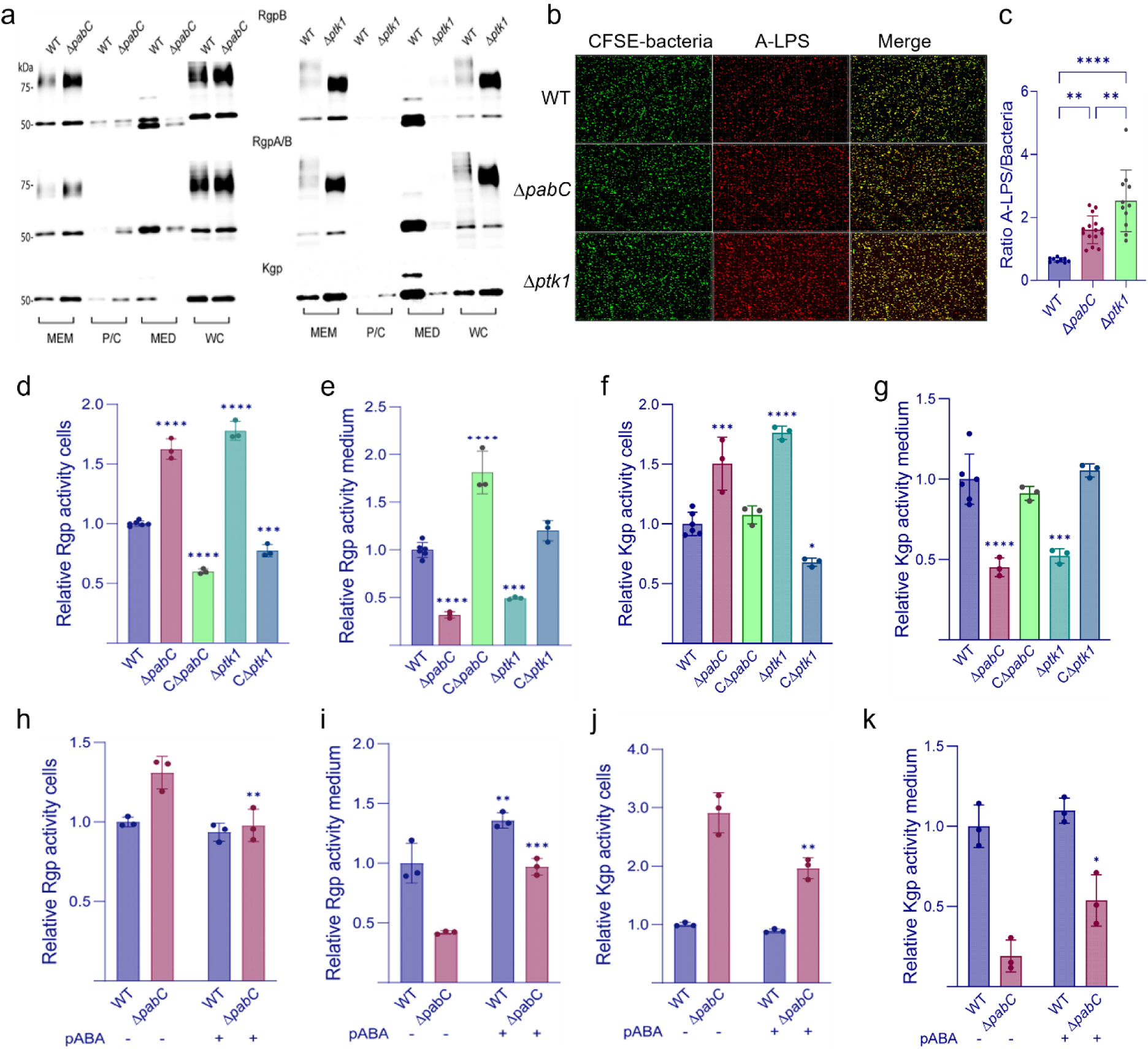
Gingipains are differentially sorted in mutants lacking PabC or Ptk1. a) Western blots of cell fractions of WT, Δ*pabC* and Δ*ptk1* strains. MEM: total membrane fraction; P/C: periplasmic and cytoplasmic fraction; MED: extracellular fraction; WC: whole cell lysate. Blots were probed with antibodies recognizing RgpB, RgpA and B, or Kgp. b) Confocal microscopy of CFSE stained *P. gingivalis* strains probed with A-LPS antibodies and Alexa Fluor 647 labeled secondary antibodies. c) Plot of fluorescent intensities from (b). Data are expressed as mean ratios of A-LPS to bacterial cells +/- SD. \*\**P*<.01, \*\*\*\**P*<.001 by one-way analysis of variance (ANOVA) with Tukey’s correction for multiple comparisons. Results are representative of three independent experiments. d-g) Rgp or Kgp activity measured with L-BApNA and Tos-GPK-pNA as the substrate, respectively, in cells and in medium after removal of cells by centrifugation. h-k) Strains were cultured with pABA 0.25 mg/ml where indicated and gingipain activity measured as in (d-g). Data are means +/- SD and expressed relative to WT. \*\**P*<.01, \*\*\**P*<.005, \*\*\*\**P*<.001 relative to WT (d-g) or corresponding strain without pABA (h-k) by one-way analysis of variance (ANOVA) with Tukey’s correction for multiple comparisons. Data are representative of 3 independent experiments.

### Neutrophil interactions track with virulence

To gain a deeper understanding of *P. gingivalis* pathogenicity we utilized in vivo and ex vivo models. Both the Δ*ptk1* and Δ*pabC* mutants survived in the murine subcutaneous infection model (Fig. 3a), indicating efficient pABA scavenging by *P. gingivalis*. Moreover, both mutants incited lower levels of neutrophil accrual at the site of infection than the parental strain (Fig. 3b, c), signifying that disruption of neutrophil recruitment may be a pathogenic feature. Consistent with this, Δ*ptk1* and Δ*pabC* inhibited the chemotaxis of isolated human neutrophils toward CXCL1 in transwell assays (Fig. 3d).

**Figure 3.**
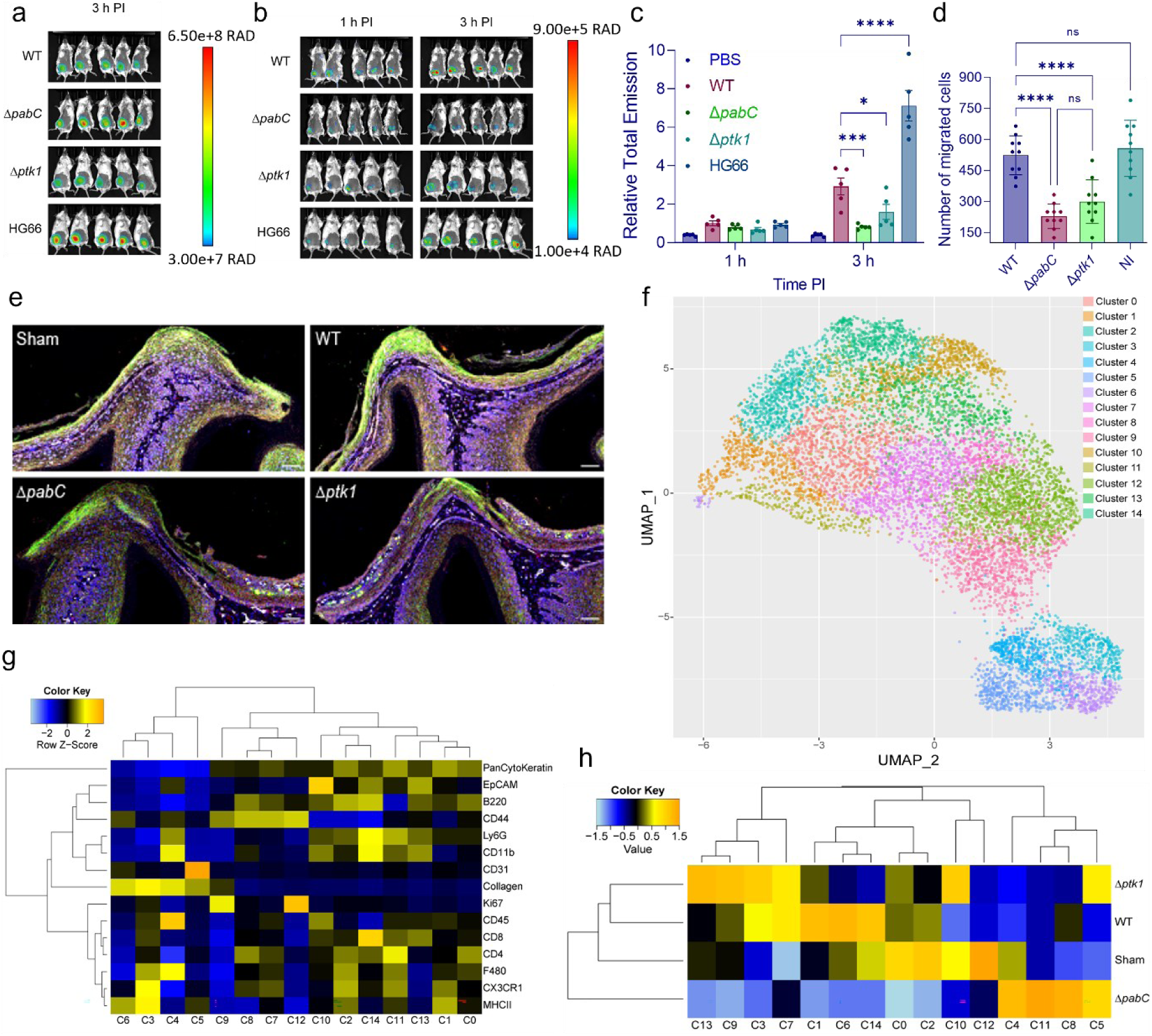
PabC and Ptk1 are required for neutrophil mobilization and recruitment. a, b) IVIS imaging of mice inoculated sc with *P. gingivalis* strains indicated. Bacteria stained with XenoLight RediJect were imaged a 3 h post infection (PPI) in (a). Mice were injected ip with luminol to detect myeloperoxidase activity of neutrophils and imaged at 1 and 3 h (PI) in (b). The scale bar shows radiance in photons per second per square centimeter per steradian (p/sec/cm^2^/sr). c) Quantitation of radiance in (b) as means +/- SEM expressed relative to WT. \**P*<.05, \*\*\**P*<.005, \*\*\*\**P*<.001 relative to WT at 3h PI by two-way repeated measures (ANOVA) with Tukey’s correction for multiple-comparison. d) Chemotaxis of neutrophils toward CXCL1 following reaction with *P. gingivalis* strains indicated. NI indicates no infection. Data are mean numbers +/- SD of migrated cells/insert from 10 fields of view. e) Representative mass cytometry images of gingival tissue from mice infected with WT, Δ*pabC,* Δ*ptk1,* or PBS (Sham) and stained with intercalator Iridium and with antibodies recognizing Ly6G (green), CD11b (red) and CD31 (white). Bar = 50 μm. f) Two-dimensional UMAP plot utilizing the expression levels of lineage markers across all tissue samples to project and visualize 15 phenotypic clusters of cells (C0-14), determined via unsupervised clustering with PhenoGraph. g) Heat map characterizing lineage marker phenotype of each cluster. h) Heat map of cell cluster representation in gingival tissue from mice infected WT, Δ*pabC,* Δ*ptk1,* or PBS (Sham).

Neutrophil recruitment to gingival tissue following oral infection of mice was further examined in vivo using imaging mass cytometry (IMC) for profiling immune/inflammatory cells in the tissue. Fig. 3e shows representative images of gingival tissue stained with antibodies reactive to Ly6G (a pan-neutrophil lineage marker) and CD11b (a murine neutrophil marker) and CD31, a cell adhesion molecule involved in neutrophil recruitment to inflamed site. Additional antibody staining is shown in Fig S4. Dimensionality reduction of all antibody staining data using Uniform Manifold Approximation and Projection (UMAP) into two dimensions was used as the basis to project 15 phenotypic groups of cells using unsupervised clustering via PhenoGraph (Fig. 3f). The phenotypic makeup of each cluster allowed us to compare the relative expression of cell marker phenotypes present in gingival tissue from mice infected with different *P. gingivalis* strains. Cluster (C) 14 is characterized by higher expression of Ly6G+ and CD11b+, while strong expression of CD31 was unique to C5 (Fig. 3g). The heat map in Fig. 3h shows that C14 is reduced in the Δ*pabC* and Δ*ptk1* conditions compared to WT and Sham, consistent with reduced neutrophil recruitment in response to these strains. In contrast, C5 overrepresentation was associated with the mutant strains, indicating a compensatory response by the host to enhance neutrophil recruitment to the site of infection. Interestingly, Δ*pabC* and Δ*ptk1* distinctly occupy separate subclusters on the heatmap, indicating differences in other cell populations, including macrophages, B-cells, and T-cells. Hence, these cell types may play a less significant role in *P. gingivalis* virulence compared to neutrophils.

Gingipains have been documented to impinge on neutrophil phagocytosis, intracellular killing and NET formation (23),(24). Thus, we dissected the outcomes of human neutrophil interactions with the mutant strains. Both Δ*pabC* and Δ*ptk1* were opsonized more efficiently than WT by IgM, although not by IgG or C3b (Fig. 4a, fig. S5a, b) and were phagocytosed more efficiently compared to WT (Fig. 4b). This may be related to another independent function of Ptk1, that of contributing to extracellular polysaccharide (EPS) production. We have previously established reduced EPS production in the *ptk1* mutant of strain 33277 (14). Similarly, fluorescent staining of EPS production by Δ*pabC* showed diminished levels, consistent with a role for PabC in regulating Ptk1 activity (Fig. S5c). Despite increased phagocytosis however, both mutants were more resistant to killing by human neutrophils (Fig. 4c), and stimulated lower levels of extracellular and intracellular ROS production (Fig. 4d). Consistent with this, colocalization of myeloperoxidase with Δ*pabC*- and Δ*ptk1*-containing phagosomes was diminished (Fig. 4e). Production of neutrophil extracellular traps (NETs) was also reduced by both mutants (Fig. 4f), and Δ*pabC*, but not Δ*ptk1,* also induced less elastase degranulation (Fig. 4g). Extracellular lactoferrin and MMP-9 levels were unaffected by both mutants (Fig. S5d, e), signifying no change in exocytosis of specific granules or gelatinase granules. These results suggest a pattern whereby reduced EPS leads to an increase in opsonization and phagocytosis; however, retention of active gingipains on the cell surface confers resistance to phagosome maturation and neutrophil killing mechanisms. We thus concluded that surface gingipains make a greater contribution to virulence compared to soluble enzymes released into the external milieu. This was further corroborated using *P. gingivalis* strain HG66 a naturally occurring mutant in *wbpB* which cannot make A-LPS and thus is unable to retain gingipains on the cell surface (25). Genomically, HG66 is very similar to 33277 (26), with an average nucleotide identity of 99.74% (https://github.com/ParBLiSS/FastANI), but colonization does not corelate with disease activity (27). Compared to WT, strain HG66 was more susceptible to neutrophil killing (Fig. 4c) and in the mouse model, HG66 incited significantly more neutrophil mobilization (Fig. 3b, c).

**Figure 4.**
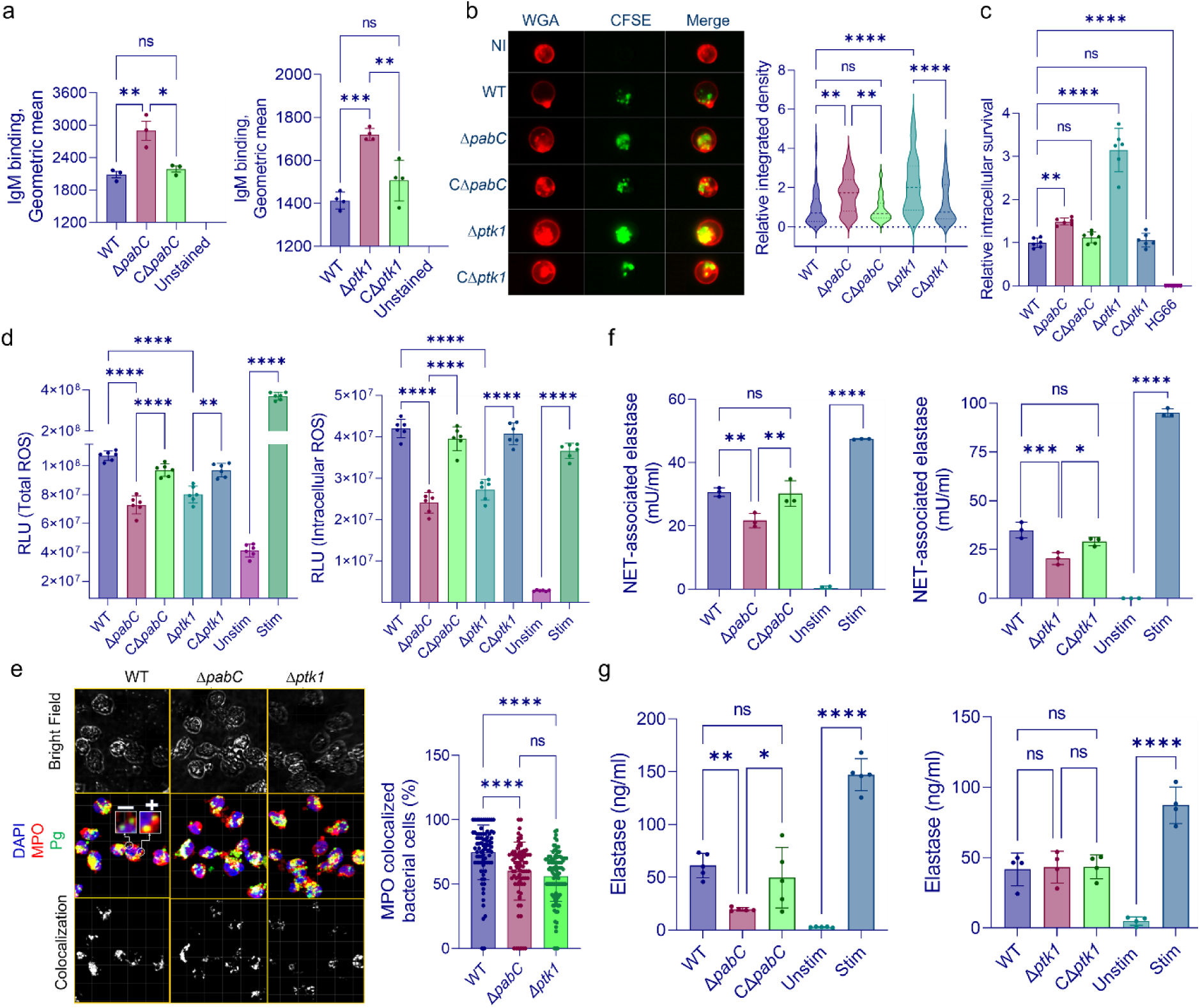
Role of PabC and Ptk1 in resistance to human neutrophil functions. a) Opsonization of *P. gingivalis* strains is indicated in 10% human serum. Bacteria were probed with allophycocyanin (APC) conjugated IgM and analyzed by flow cytometry. Data are geometric means ± SD of fluorescence. b) Left panel is an Amnis Imagestream representative image of phagocytosis of *P. gingivalis* strains. Alexa Fluor 647 conjugated WGA-stained neutrophils (red) were reacted for 30 min with serum-opsonized CFSE-stained *P. gingivalis* strains (green). NI indicates no infection. Right panel is a plot of CFSE fluorescence associated with neutrophils from at least 40 cells. Values are integrated density relative to WT. c) Intracellular survival of serum opsonized *P. gingivalis* strains after incubation with neutrophils at MOI 10 for 30 min. Values represent CFU cultured postlysis and are means +/- SD expressed relative to WT. d) Total (left panel) or intracellular (right panel) ROS production by neutrophils following challenge with serum-opsonized *P. gingivalis* strains at MOI 10 for 1 h. ROS was measured by luminol chemiluminescence and data are means +/- SD. Unstim is unstimulated negative control and Stim is positive control with PMA. e) Left panel is confocal image of neutrophils challenged with CFSE-labeled *P. gingivalis* strains (MOI 10) and reacted with antibodies to MPO (red). Boxes represent examples of colocalization (+) or separate bacteria and MPO reactivity (-). Right panel is quantitation of image data from 80 cells and are means +/- SD. f) NET formation by neutrophils challenged with *P. gingivalis* strains MOI 10 for 4h. Data are means +/- SD. Unstim is unstimulated negative control and Stim is positive control with PMA. g) Elastase degranulation. Neutrophils challenged with serum-opsonized *P. gingivalis* strains MOI 10 for 4 h. Data are means +/- SD. Unstim is unstimulated negative control and Stim is positive control with latrunculin and fMLF. \**P*<.05, \*\**P*<.01, \*\*\**P*<.005, \*\*\*\**P*<.001 by ANOVA with Tukey’s correction for multiple comparisons. ns is not significant (*P*>.05). Data are representative of at least three independent experiments.

### Role of Ptk1 in One Carbon Metabolism

Our results show that levels of pABA influence Ptk1 activity, which in turn defines pathogenic potential. We next sought to determine if Ptk1 can reciprocally regulate OCM. Folate levels are a measure of OCM flux (28, 29), and Δ*pabC*, which lacks the essential OCM precursor pABA, was deficient in folate production, a deficiency which was reversed by the addition of exogenous pABA (Fig. 5a). The gingipain-null mutant also exhibited lower folate levels, establishing the importance of a source of exogenous amino acids for flux through OCM in *P. gingivalis*. Additionally, Δ*ptk1* was deficient in the biosynthesis of folate, suggesting that soluble gingipains are more effective than membrane associated at delivering peptides necessary for further intracellular processing to amino acids (30) that serve as substrates for OCM (31). This conclusion is supported by the finding that strain HG66 produced more folate than the WT strain. Another possible role for Ptk1 is in the direct regulation of OCM enzyme activity. To address the second possibility, FLAG-tag Ptk1 was expressed in *P. gingivalis* and after immunoprecipitation with FLAG antibodies, potential substrates were identified by MS/MS. Supplementary Data S3 shows that among the binding partners of Ptk1 were 4 enzymes involved in OCM, namely GcvT aminomethyltransferase (*gcvT,* PGN_0550), GlyA serine hydroxymethyltransferase (PGN_0038), ALP alkaline phosphatase (PGN_1049) and FolP dihydropteroate synthase (PGN_0522). Additionally, Rgp and Kgp peptides were identified, consistent with the ability of Ptk1 to directly phosphorylate gingipains. A phosphorylation assay with purified Ptk1 and substrate proteins confirmed that GcvT, GlyA and ALP can be phosphorylated by Ptk1, while phosphorylation of FolP could not be detected in this *in vitro* assay (Fig. 5b). We also demonstrated that the enzyme activity of ALP was increased against pNPP substrate following phosphorylation by Ptk1 (Fig. 5c). ALP functions in the pathway leading to 6-hydroxymethyl-7,8-dihydropteridine-P2 (DHPPP) which combines with pABA to form DHP, catalyzed by FolP. Thus, a reduction in Ptk1-dependent phosphorylation will result in lower levels of DHP and less flux through OCM. Moreover, in OCM one-carbon units are predominantly derived from serine and glycine, which can be synthesized *de novo* or obtained exogenously (31). As both GcvT and GlyA are involved in the metabolism of serine and glycine, Ptk1 can exert multi-level control over the availability of one-carbon units. pABA in turn provides positive feedback from OCM to Ptk1 through inhibition of the Ltp1 phosphatase.

**Figure 5.**
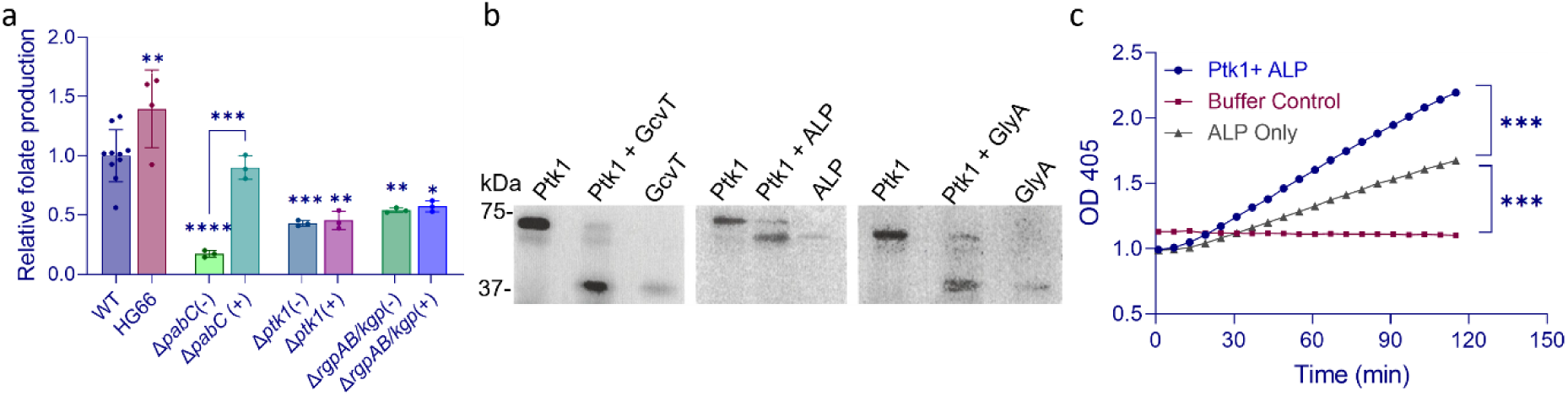
Effects of PabC and Ptk1 on OCM. a) Folate production by strains of *P. gingivalis*. Folate released from *P. gingivalis* cell lysates was added to a folate auxotroph of *L. rhamnosus* and OD_600_ measured after growth for 18 h. *P. gingivalis* strains were either cultured with (+) or without (-) pABA 0.25 mg/ml. Data are means +/- SD and are expressed relative to WT. \**P*<.05, \*\**P*<.01, \*\*\**P*<.005 by ANOVA with Tukey’s correction for multiple comparisons, relative to WT except where indicated. pABA had no significant effect (*P*>.05) on folate production by Δ*ptk1* or Δ*rgpAB/kgp*. Data are representative of three independent experiments. b) Substrate phosphorylation by Ptk1. Recombinant substrate proteins (GcvT: 40 kDa; ALP: 62 kDa; GlyA: 42 kDa) were dephosphorylated with CIP and reacted with the catalytic domain of Ptk1 in the presence of ^32^P labelled ATP for 1 h. The reaction was separated by SDS-PAGE and phosphorylation visualized by autoradiography. Image is representative of three independent experiments. c) Enzyme reaction of ALP following phosphorylation by the catalytic domain of Ptk1. Phosphorylated or control ALP was reacted in triplicate with pNPP and OD_405_ measured over time. \*\*\**P*<.005 by Student’s t-test. Data are representative of three independent experiments.

## Discussion

At mucosal barriers the complex and dynamic nutrient pool requires that microbial colonizers implement adaptable regulatory networks to maintain optimal physiology. The bacterial tyrosine (BY) kinase family of enzymes integrates into complex signaling networks, which can regulate virulence-associated properties including polysaccharide capsule biogenesis and secretion of T3SS proteins (32–34). BY kinase functionality is driven by the opposing actions of intermolecular autophosphorylation and activation, which is counteracted by dephosphorylation and inactivation catalyzed by tyrosine phosphatases (35, 36). Here, we uncover a BY kinase-dependent physiology-virulence coupling mechanism in the oral pathobiont *P. gingivalis*. The OCM precursor pABA regulates the activation state of the BY kinase Ptk1 through inhibition of the phosphatase Ltp1 (13), and in the absence of pABA, Ptk1 balance shifts toward an inactive state. Inactivation, or loss, of Ptk1 drove an increase in virulence in animal models which was associated with retention of gingipain proteases in the cell membranes and a conjoint reduction in secreted gingipains.

The gingipain proteases are multifunctional effector molecules which are delivered to the outer membrane through the type IX secretion system (T9SS) (37, 38) and tethered to the cell surface following PorU-catalysed attachment of anionic (A)-LPS to the C-terminus by transpeptidation (39). Amino groups of exogenous amino acids and peptides can compete with the NH_2_ group on A-LPS and release gingipains into the extracellular milieu. Soluble gingipains are also released when the transient thioester bond with the PorU sortase is hydrolyzed. As an inner membrane protein, Ptk1 is positioned to interact with gingipains as they enter the secretion apparatus. Loss or suppression of Ptk1 resulted in retention of gingipains in the membrane fraction, indicating that gingipain phosphorylation is required for either sequential movement through the T9SS machinery, or for disassociation from the membrane. The finding that the gingipains are fully processed to the membrane-type isoforms anchored in the outer membrane suggests that transport through the secretion machinery and proteolytic processing occur as normal in the Δ*pabC* and Δ*ptk1* mutants, and that phosphorylation may facilitate hydrolysis of the PorU-gingipain thioester bond. Indeed, in other systems, phosphorylation can enhance access to thioester enzymes (40), and increase the rate of hydrolysis of thioester bonds (41).

While it is long established that gingipains can localize to both the cell surface and extracellular milieu, the relative contribution of these fractions to virulence has been a matter of speculation. One widespread view is that extracellular enzymes target host tissue and immune effector molecules more broadly as well as provide blanket protection to non-proteolytic pathobionts. Here we demonstrate the converse situation whereby an increase in surface gingipain activity along with a decrease in soluble gingipain correlate with reduced neutrophil mobilization and elevated pathogenic potential in animal models of disease. Thus, soluble gingipains may represent important PAMPs that evoke protective innate immune responses. Moreover, we found that surface retention of gingipains was associated with resistance to neutrophil killing mechanisms and enhanced survival. In contrast, opsonization by IgM and neutrophil phagocytosis were increased in the mutant strains, although this may reflect the reduction of EPS when Ptk1 activity is suppressed. Hence, *P. gingivalis* virulence is tunable by Ptk1, with more virulent states capable of suppressing recruitment and surviving the antibacterial mechanisms deployed by neutrophils. The stimuli that initiate activation of BY kinases in general are not well-defined and it is likely that the Ptk1 signaling node may be a target for other host or microbiome factors that influence the pathogenic state of *P. gingivalis*.

Both *ptk1* and *pabC* mutants exhibited similar pathogenic potential, consistent with a role for pABA in activating Ptk1. However, inhibition of Ptk1 was only partial in the absence of pABA and hence pABA may have additional functionality in the control of the *P. gingivalis* pathogenic state. For example, OCM metabolites regulate the fidelity and rate of translation initiation (5). Additionally, the alarmone ZTP senses insufficient stores of intracellular folate and coordinates adaptive responses to compensate and restore folate to sufficient levels (42). pABA levels and OCM may, therefore, have a broad impact on the overall physiology of *P. gingivalis*. Moreover, RNA-Seq analysis indicated loss of pABA may increase CRISPR activity. Nevertheless, posttranscriptional regulation was of paramount importance in determining *P. gingivalis* pathogenic state and reliance on the transcriptional landscape alone would not have provided a sufficient understanding of the process.

Flux through OCM depends on substrate concentration and on the number of one-carbon units available for transfer (43, 44). Ptk1 regulated both processes through phosphorylation of gingipains and OCM metabolic enzymes. It is tempting to speculate, therefore, that Ptk1 can maintain a balance of one-carbon unit availability through regulating both gingipain-dependent generation of serine and glycine from exogenous proteins as well as the metabolic enzymes that mediate their biosynthesis/degradation. Moreover, Ptk1 can modulate the pathway leading to the production of DHPPP which is subsequently combined with pABA. Balance in the OCM pathway can then be preserved through pABA-dependent inactivation of Ltp1 and activation of Ptk1. In this manner Ptk1 metabolically couples OCM with gingipain location-dependent virulence; however, the increase in virulence can be seen as an ‘inadvertent’ consequence of the action of gingipains on host tissues and neutrophil function. *P. gingivalis* is an inhabitant of the oral microbiome in health and in disease (45) and is adapted to human ecosystems. In contrast to other organisms, the pABA auxotroph of *P. gingivalis* can survive in vivo, and *P. gingivalis* can scavenge exogenous pABA from microbiome partner species such as *S. gordonii*. Sufficient pABA helps maintain *P. gingivalis* in a non-pathogenic state (9); however, this calming influence will be lost when pABA production by partner organisms is reduced or communities become over-represented with pABA consuming organisms. For example, when *P. gingivalis* and *S. gordonii* cells become physically integrated, pABA production by *S. gordonii* is suppressed and virulence is enhanced (9, 46). Other periodontal organisms with which *P. gingivalis* can associate, including *Fusobacterium nucleatum, Tannerella forsythia* and *Filifactor alocis*, lack obvious homologs of the genetic machinery to produce pABA *de novo* and are thus likely net pABA consumers, which will relieve pathogenic constraints on *P. gingivalis*. Synergistic interactions between *P. gingivalis* and *T. denticola* may also involve OCM-based responses as treponemal conditioned growth medium differentially regulated OCM pathway genes in *P. gingivalis* (47). Collectively, these results provide novel insights into the mechanisms that control the pathogenic potential of endogenous microbiome constituents and help to resolve the longstanding paradox regarding *P. gingivalis* which, despite possessing a suite of potent virulence factors (48), remains non-pathogenic in most instances.

## Methods

### Bacteria, mutant construction and growth conditions

*P. gingivalis* 33277 (WT), HG66, and isogenic mutants, were cultured anaerobically (85% N_2_ 10% CO_2_, 5% H_2_) at 37^0^C in Trypticase soy broth (TSB) supplemented with 1 mg/ml yeast extract, 5 μg/ml hemin, and 1 μg/ml menadione. When necessary, erythromycin (10 μg/ml), tetracycline (1 μg/ml), or chloramphenicol (20 μg/ml) was added to the medium for isogenic mutants. *Escherichia coli* S17-1, BL21 Star (DE3) were grown in LB broth aerobically at 37^0^C with shaking, and containing when necessary kanamycin (50 μg/ml) and ampicillin (100 μg/ml). *Lactobacillus rhamnosus* ATCC 7469 was cultured in MRS broth at 37^0^C in 5% CO_2_.

*P. gingivalis* mutant strains Δ*ptk1,* C*Δptk1* a strain of Δ*ptk1* complemented in trans with the wild type allele, *Δptk1:ptk1*FLAG a strain of Δ*ptk1* complemented in trans with the wild type allele fused to the FLAG epitope, and Δ*rgpAB/kgp* a gingipain null mutant were reported previously (14, 49). An allelic exchange Δ*pabC* mutant was constructed as described (50). Briefly, ∼800 bp regions flanking *pabC* (PGN_1333) were fused to the *ermF* gene by commercial synthesis (Genewiz). The synthesized construct was amplified using the Phusion High-Fidelity PCR Kit (New England Biolabs) with the primers listed in table S1, and gel purified using the Monarch DNA Gel Extraction Kit (New England Biolabs). One µg of the amplified and purified construct was electroporated into electrocompetent *P. gingivalis* WT, and transformants were selected on TSB agar plates supplemented with erythromycin. The mutation was confirmed by PCR and southern blot analysis. Loss of mRNA expression from *pabC* along with undisrupted expression from downstream genes was confirmed by RT-PCR and RNASeq analysis.

To generate the CΔ*pabC* complemented strain, the promoter region 475 bp upstream of *pabB* and the coding region of *<u>pabC</u>* were synthesized (Genewiz), fused and cloned into pT-COW. The resulting plasmid, pTpabC, was transferred into the Δ*pabC* strain through conjugation with *E. coli* S17-1 as described (15). Transconjugants were selected with gentamicin, erythromycin, and tetracycline, and expression of *pabC* mRNA confirmed by RT-PCR.

### Antibodies

Rabbit antibodies to purified RgpB, which also recognize the catalytic domain of RgpA, designated RgpA/B antibodies are described previously (51). Mouse monoclonal antibodies (MAbs) specific for the Kgp catalytic domain (7B9) and to a single epitope on RgpB (18E6) were developed at the University of Georgia Monoclonal Antibody Facility using recombinant protein as the antigen (52, 53). *P. gingivalis* anti-anionic (A)-LPS (1B5) MAb was a kind gift from Dr. Mike Curtis (St. Bartholomew’s & the Royal London Queen Mary’s School of Medicine & Dentistry, UK).

### Bacterial fractions

*Porphyromonas gingivalis* strains were grown to mid-log phase and cultures were adjusted to OD_600_ = 0.8. One mM tosyl-L-lysyl-chloromethane hydrochloride (TLCK), Halt Protease and Phosphatase Inhibitor Cocktail (ThermoFisher), and Phosphatase Inhibitor Cocktail (ThermoFisher) were added. Cells were collected by centrifugation for 15 min at 6000 g and 4^0^C, and the supernatant collected as the medium fraction. The pellet was washed twice with phosphate-buffered saline (PBS), resuspended in the initial volume of PBS with added inhibitors, and disrupted with BugBuster buffer (MilliporeSigma) to prepare the cell lysate. For periplasm/cytoplasm preparation, pellets resuspended in PBS were sonicated on ice with 6 x 5 s pulses (17 W per pulse) and a 10 s rest between each pulse. The lysate was centrifuged at 1000 g for 15 min at 4^0^C, and the supernatant collected and centrifuged at 150,000 g for1 h at 4^0^C. The resulting supernatant was designated as the periplasm/cytoplasm fraction. The pellet was resuspended in 5 ml PBS, dispersed by sonication with 3x 5 s pulses in an ice-water-bath and designated the total membrane fraction.

### Mice

Male and female germ-free (GF) BALB/c mice (Taconic Biosciences) were housed in isolators at the University of Louisville Germ-free Animal Research Facility. Littermates were randomly assigned to experimental groups. The sterility of GF animals was confirmed by aerobic and anaerobic culture of oral swabs and fecal pellets on non-selective media, and by PCR using universal 16S primers. Conventional mice (BALB/c, Jackson Laboratories) were housed in the animal care facilities of the University of Louisville.

### Murine periodontitis model

Conventional or GF mice of both sexes, 6–10 weeks old, were orally inoculated using a ball-end needle 5 times at 2 days intervals with *P. gingivalis* WT or isogenic mutant strains (10^9^ CFU) suspended in 0.1 ml sterile PBS with 2% carboxy-methylcellulose (CMC). PBS with CMC alone was used for sham inoculations. Forty-two days after the last infection, mice were euthanized and skulls were subjected to micro-computed tomography (μCT) scanning (SkyScan 1174; Bruker). Bone loss was assessed by measuring the distance between the alveolar bone crest and the cementoenamel junction at 6 interdental points at 3 sites around the first and second maxillary molars.

### Imaging Mass Cytometry

Conventional mice were inoculated with *P. gingivalis* strains as above, and after euthanization the gingival tissue surrounding the three maxillary molars was removed. The tissue was fixed in 10% neutral-buffered formalin at 4^0^C overnight, followed by three washes with PBS at room temperature (RT) for 5 min each. The fixed gingival tissue was dehydrated using graded ethanol and chloroform and then embedded in paraffin. Tissue blocks were sectioned to a thickness of 8 µm, mounted on slides, and dried at 37°C overnight. IMC staining was performed according to the Standard BioTools protocol. Briefly, paraffin embedded mouse gingival sections were deparaffinized with Xylene (5 min, 3 times) following by re-hydration for 5 min in descending grades of ethanol (100%, 95%, 80%, 70%). The final re-hydration was in Milli-Q water for 5 min. Antigen retrieval was performed in a Steamer with pre-heated Antigen retrieval buffer for 30 min, and after cooling to RT samples were washed with Milli-Q water and blocked with 3% BSA in DPBS (Dulbecco′s Phosphate Buffered Saline, SigmaAldrich) for 45 min at room temperature. Primary antibodies were purchased in ready to use format from Standard BioTools (table S2), added to DPBS containing 0.5% BSA, and reacted with sections overnight at 4°C. The slides were washed 2 x 5 min in DPBS containing 0.2% Triton X-100 followed by 2 x 5 min washes in DPBS. Tissue was stained with 125 µM Intercalator Iridium in DPBS at 1:400 for 30 min at RT. The slides were washed for 5 min in DPBS followed by a quick rinse with Milli-Q water. Slides were dried at RT and stored in a dust-free and dry container at 4^0^C until laser ablation by the Hyperion Imaging System (Standard BioTools). 400 µm X 800 µm of the region of interest (ROI) per sample were ablated. MCD-files created by the CyTOF® Software v7.0 were processed in MCD viewer (Standard BioTools) to create pseudo-colored images for marker expression visualization. Software of Ilastic, CellProfiler and HistoCat for image analysis were used for cell segmentation, image masks and fcs file conversion. Further downstream analyses utilized *PICAFlow*, a workflow written in R for flow and mass cytometry analysis (54). In brief, fcs files were used as input and underwent pre-processing steps to rename and subset the file parameters to include only cell marker signal data. Further processing was via arcsinh transformation of the data with subsequent normalization to correct for potential batch and erroneous inter-sample heterogeneity effects by alignment of signal peaks across all samples. The transformed and normalized signal data were used to test for optimal settings for UMAP dimensionality reduction which was determined to be 0.3 for the minimum distance value and 391 for number of neighbors value. These final settings were used to apply UMAP dimensionality reduction on the entire signal dataset and generate a 2-dimensional plot of all cells in the dataset. Next, clustering analysis of the signal data was performed first by testing the normality of the signal data for each cell marker, and the result showed non-normal results for all markers, so the median was used as the metric for further cluster analyses instead of the mean. Next, the PhenoGraph method (55) as implemented in *PICAflow* was used with a k of 100 to perform the clustering of cells using the signal data and the results were projected onto the previously generated UMAP plot. The clustering data was further used to generate heatmaps to visualize the relative expression of cell markers within each cluster and the relative amount of cells belonging to each cluster per treatment group.

### Murine neutrophil MPO activity

Conventional BALB/c mice, 8-10 weeks-old, were inoculated subcutaneously (sc) with 0.1 ml of 3 x 10^9^ *P. gingivalis* cells suspended in sterile, prereduced PBS, or with PBS alone as a control. Prior to inoculation, the hair on the abdomen and inner thigh of each mouse was removed using Nair. At 1 and 3 h post inoculation, mice were placed under anesthesia using isoflurane and each mouse was injected intra-peritoneally with 300 mg/kg of bodyweight of 50 mg/ml filter sterilized luminol sodium salt solution (MilliporeSigma). After 5 min incubation, *in vivo* chemiluminescence imaging was performed at 120 s exposure using a Spectral Ami HT (Spectral Instruments Imaging). For bacterial detection, bacteria were pre-labelled with 200 XenoLight RediJect Bacterial Detection Probe (Perkin Elmer) for 15 min at RT, then washed 3 times with sterile, pre-reduced PBS. Bacterial fluorescence was detected in mice 3 h post infection at 1 sec exposure. Image validation, acquisition, and data analysis were performed with Aura imaging software.

### Proteolytic activity

The amidolytic activity of arginine-specific (Rgp) and lysine-specific (Kgp) gingipains was assayed as described previously (9). When necessary, cultures were supplemented with pABA (0.25 mg/ml) or vehicle control. Cells were separated from the culture supernatant fraction by centrifugation (5,000 g, 20 min), washed and resuspended in PBS to the original volume. The chromogenic *p*-nitroanillide substrates N-Benzoyl-L-Arginine-*p*NA (L-BApNA) or toluenesulphonyl-glycyl-prolyl-L-lysine *p*NA (Tos-GPK-pNA) (MilliporeSigma) were used to detect RgpA/B and Kgp, respectively. Samples (50 μl) were preincubated in 200 mM Tris HCl, 5 mM CaCl_2_, 150 mM NaCl_2_, 10 mM cysteine in 96-well plates for 10 min at 37^0^C and assayed with 0.5 mM substrate. The rate of substrate hydrolysis and the accumulation of *p*-nitroanilide were monitored spectrophotometrically at 405 nm over time in a SpectraMax Plus 384 (Molecular Devices), and the activity of the enzyme expressed relative to WT.

### RNA-Seq and data analysis

Mice were inoculated sc as described above. Skin abscesses were allowed to develop for 48 h and abscess material was collected using sterile cotton swabs and suspended in sterile, prereduced PBS at 4°C. Samples were centrifuged at 150 g for 3 min at 4°C to pellet any residual mouse cells and the supernatant collected and centrifuged at 5000 g for 5 min at 4°C to recover the bacterial cells. The pelleted bacteria cells were resuspended with TRIzol™ Reagent (Invitrogen) and total RNA extraction was caried out using the Qiagen RNeasy Plus Mini Kit (Qiagen). RNA sequencing using 150 bp paired-end reads was performed by a commercial vendor (Novogene) using an Illumina NovaSeq 6000. The quality of reads and trimming was with *fastP* (56). The trimmed reads were aligned to a combined *Mus musculus* (GRCm39 Primary Assembly) and *P. gingivalis* ATCC 33277 genome (accession AP009380.1) using the *Hisat2* aligner (57) to generate SAM files that were subsequently converted to coordinate-sorted BAM files. Reads aligning only to *P. gingivalis* were split into separate BAM files. Raw counts were generated by parsing the *P. gingivalis*-only BAM files using *featureCounts* (58). A count for a gene was considered valid if a read overlapped at least 7 base pairs with the gene. Any counts remaining from deleted genes in the corresponding deletion mutant strains were manually set to 0 after inspecting the alignment and ensuring no genuine reads mapped to these deleted genomic regions. The raw counts per gene were input into the *DESeq2* Bioconductor/R package (59) using the recommended guidelines, including filtering out genes with total counts less than 10 across all samples. Output included Log2 fold change expression values and *P*-values adjusted for multiple comparisons using the Benjamini-Hochberg procedure (60). Log fold-change shrinkage was applied to the output via *apeglm* (61) as implemented within *DESeq2* to reduce the false-positive rate and refine the ranking of genes. This output was used as input for the generation of volcano plots using the *EnhancedVolcano* Bioconductor/R package (https://github.com/kevinblighe/EnhancedVolcano), and for functional enrichment analysis through the STRING Database version 12.0 (62) using an FDR stringency of 25 percent and a minimum interaction confidence score of 0.4 for network generation. For principal component analysis (PCA) and heatmap generation, the raw count data were made homoscedastic using a regularized logarithm transformation (59). PCA plots were generated using base R, *ggplot2* (63), and *ggforce* (64) packages for R. Gene count data for heatmap generation were further converted into z-scores and used as input into the *ComplexHeatmap* Bioconductor/R package (65). Visualization of sets of genes in common amongst strains was performed using Venn diagrams generated through the *ggvenn* R (https://github.com/yanlinlin82/ggvenn).

### Mass spectrometry

Samples were separated by SDS-PAGE and lanes excised for in-gel trypsin proteolysis as described previously (66). The concentrated and desalted peptides were re-suspended in 20 μl of 2% acetonitrile/0.1% formic acid, and filtered through a 0.45 μm regenerated cellulosic filter, prior to analysis by 1-dimensional (1D) reversed phase liquid chromatography (RP-LC) mass spectrometric analysis (MS). An equal volume of tryptic peptides from each digest was separated using an EASY n-LC (ThermoFisher) UHPLC system and an Acclaim PepMap 100 75µm x 2cm, nanoViper (C18, 3 µm, 100 Å) trap coupled to an Acclaim PepMap RSLC 50 µm x 15 cm, nanoViper (C18, 2 µm, 100 Å) separating column (ThermoFisher). Following injection of the sample onto the column, separation was accomplished with a 50 min linear gradient from 2% to 37% acetonitrile in 0.1% formic acid.

The eluate was introduced into the LTQ-Orbitrap ELITE + ETD mass spectrometer using a Nanospray Flex source (ThermoElectron). A Nth Order Double Play method was created in Xcalibur v2.2. Scan event one of the method obtained an FTMS MS1 scan (normal mass range; 120,000 resolution, full scan type, positive polarity, profile data type) for the range 300–2000m/z. Scan event two obtained ITMS MS2 scans (CID activation type, normal mass range, rapid scan rate, centroid data type) on up to twenty peaks that had a minimum signal threshold of 5,000 counts from scan event one. The lock mass option was enabled (0% lock mass abundance) using the 371.101236m/z polysiloxane peak as an internal calibrant.

Peaks Studio Xpro (Bioinformatics Solutions) was used to analyze the data collected by the mass spectrometer. The database used in the Peaks searches was the 9/30/2021 version of the *P. gingivalis* strain ATCC 33277 protein sequences from the UniProtKB (proteome identifier UP000008842). The enzyme selected was semi specific Trypsin (maximum two missed cleavages). Fragment tolerance was 0.5 Da and parent tolerance was 15 ppm (monoisotopic). FDR estimation was enabled in the algorithms. A proteins.csv file was exported for curation in Microsoft Excel. Proteins and peptides were accepted at the 1% FDR threshold.

### A-LPS assay

*P. gingivalis* cells were washed twice with PBS at 5,000 g for 5 min, and stained with CFSE (40 μg/ml in DMSO) for 30 min at RT in the dark. After washing twice with PBS, 1×10^7^ bacteria were centrifuged (300 g, 3 min) on to glass slides and fixed with 4% paraformaldehyde for 10 min. Slides were washed twice with PBS, blocked in 5% goat serum for 1 h and reacted with A-LPS primary antibody 1:1000 overnight at 4°C. Secondary antibody was Alexa Fluor 647 labelled goat α-mouse IgG (Invitrogen) 1:500 and was incubated for 1 h in the dark at RT. Slides were viewed by laser scanning confocal microscopy (Leica, SP8). Images were acquired and analyzed using Volocity 6.3 (PerkinElmer). Ten independent images were analyzed for each condition.

### EPS staining

*P. gingivalis* cells were washed with PBS at 5,000 g for 5 min, and labelled with 10 μM Syto 17 (Molecular Probes) for 30 min at 4°C with rocking in the dark. Cells were washed once in 1x prereduced PBS and re-suspended in TSB:PBS 1:1. 3 x10^8^ *P. gingivalis* cells (500 μl) were deposited on a glass chambered slide and incubated anaerobically at 37°C for 24 h. The supernatant was removed and cells were fixed in 4% paraformaldehyde with 10 mM cetylpyridinium chloride for 15 min at RT. Cells were washed 3 times with PBS and treated with 188 nM Concavalin A-FITC and 286 nM WGA-FITC (both from Sigma) in 1ml PBS for 30 min in the dark on a rocking platform at RT. After washing three times with PBS, cells were visualized with a Leica SP-8 confocal microscope. Image analysis and intensity measurements were performed using Volocity software.

### Recombinant protein expression

To enhance the production and stability of the recombinant Ptk1 catalytic domain, pREP4-*groESL* (kindly provided by Dr. Christophe Grangeasse (67) was transformed into *E. coli* BL21 Star (DE3) cells containing the pGEX-4T-fPtk1 (including the C-terminal catalytic domain, aa 541-821 (14)) and maintained in LB broth containing kanamycin and ampicillin. Glutathione S-transferase (GST)-tagged Ptk1 expression was induced with 100 µM IPTG and purified using glutathione resin (GenScript). Soluble Ptk1 was dialyzed against Tris-buffered saline (TBS, pH 7.4), and the purity was assessed by SDS-PAGE and Coomassie staining.

Recombinant His-Tag GcvT (PGN_0550), GlyA (PGN_0038), Dihydropteroate synthase (PGN_0552), and ALP (PGN_1049) were produced by a commercial vendor (Genscript). Amino acid sequences were codon optimized for expression in *E. coli*. Synthesized constructs were cloned into pET30a, using sites NdeI (CATATG) and HindIII (AAGCTT) and transformed into *E. coli*. More details on purification method, concentration and purity achieved, and storage buffer composition are in table S3

### Quantitative RT-PCR

Total RNA was isolated from bacterial cells using an RNAeasy plus Mini kit (Qiagen) with DNase treatment, and 2 μg converted into cDNA and using LunaScript RT SuperMix Kit (Applied Biosystems). Quantitative PCR was performed by a QuantStudio 3 using the Powerup SYBR Green master mix (Applied Biosystems). Relative mRNA expression was evaluated using the 2^−ΔΔCt^ method, with 16S as the internal control. Primers were produced by Integrated Data Technologies and are described in table S1.

### SDS-PAGE and western blotting

Samples were boiled in SDS-PAGE sample buffer with 2% β-mercaptoethanol for 5 min and proteins separated by electrophoresis on SDS-PAGE gels (10% unless otherwise noted). Proteins were visualized with Coomassie Brilliant Blue (BioRad) or silver stain (ThermoFisher). For immunoblotting, proteins were electro-transferred onto nitrocellulose membranes (unless otherwise noted) and blocked with 5% non-fat milk. The primary antibodies mouse α-RgpB (1:1000), rabbit α-RgpA/B (1:10,000), or mouse α-Kgp (1:1000) were reacted for 18 h at 4^0^C. Goat α-mouse or goat α-rabbit secondary antibodies (Invitrogen) at 1:2000 were incubated for 1 h at RT. After reacting with SuperSignal West Pico Plus Chemiluminescent substrate (ThermoFisher), blots were visualized by chemiluminescent imaging (Azure Biosystems)

### Immunoprecipitation

For Ptk1 phosphorylation, *P. gingivalis* cells (2 x 10^9^) were lysed with BugBuster buffer (500 μl) containing 1 mM TLCK, Halt Protease and Phosphatase Inhibitor Cocktail, and Phosphatase Inhibitor Cocktail, followed by centrifugation at 12,000 g for 10 min at 4^0^C to remove debris. Protein concentrations were determined by a BCA assay (ThermoFisher). Proteins (150 µg) were reacted with FLAG antibodies 1:200 by rotating overnight at 4^0^C. Protein A agarose beads (ThermoFisher), washed twice in PBS, were added and reacted by rotating for 3 h at 4 ^0^C. Beads were centrifuged (1000 g, 15 s), washed twice in PBS, resuspended in 30 μl 3x Blue Sample Loading buffer (Invitrogen) with 2% β-mercaptoethanol, and boiled for 5 min. The samples were analyzed by SDS-PAGE and immunoblotting using mouse phospho-tyrosine antibody (1:1000, MilliporeSigma) and goat anti-mouse secondary antibody, 1:2,000.

For identification of Ptk1 client proteins, *P. gingivalis* cells (6 x 10^10^) were centrifuged (5,000 g, 10 min) and resuspended in 5 mL of lysis buffer composed of 50 mM Tris pH 8, 150 mM NaCl, 5 mM MgCl_2_, 5 mM TLCK (Sigma), 2% Triton X-100, 5 µL benzonase (Sigma), and Halt protease and phosphatase inhibitor. Cells were sonicated on ice with 6x 5 s pulses (17 W per pulse) with a 10 s rest between each pulse. Cellular debris was centrifuged at 10,000 g for 25 min at 4°C. Twenty µL (packed volume) of anti-FLAG magnetic beads (MilliporeSigma) were added to the supernatant and rotated overnight at 4°C. Beads were removed with a magnetic separator and washed 4x 1 min with TBS (400 µL). Protein was eluted with 0.1 M glycine pH 3 (100 µL) for 10 min with rotation. A total of 30 µL of eluate was separated on an 8% SDS-PAGE gel, and gel slices were used for mass spectrometry.

### Substrate phosphorylation

Purified substrate candidates were dephosphorylated with CIP agarose (MilliporeSigma) according to the manufacturer’s instructions. Briefly, CIP agarose was prepared by washing 3x with 1x kinase buffer and 6 µg of substrate protein were added in kinase buffer (100 mM NaCl, 50 mM Tris-HCl pH 8, 10 mM MgCl_2_, and 1 mM DTT). Reactions were incubated for 1 h at 37°C with periodic mixing, and beads were removed by centrifugation at 10,000 g for 1 min. Kinase reactions were in kinase buffer with 2 µg of dephosphorylated substrate protein, 1 µg of purified Ptk1, phosphatase inhibitor #2 cocktail (ThermoFisher), and 5.5 µCi ^32^P. Reactions were incubated for 1 h at 37°C. The entire reaction was separated by 12% SDS-PAGE and electro-transferred to a PVDF membrane. The membrane was exposed to X-ray film or storage screens which were phospho-imaged (Azure Sapphire).

### Phosphatase assay

Purified ALP (PGN_1049, 6 µg) was incubated in kinase buffer with 5 mM ATP and 2 µg Ptk1 for 1 h at 37°C in a total of 50 µL. An equal volume of 100 mM pNPP in 2x phosphatase buffer (200 mM NaCl, 100 mM Tris pH 8, 1.6 mM DTT, and 5 mM MnCl_2_) was added to each phosphatase reaction mixture in a 96 well microplate. OD_405_ was measured every minute for 120 min (Spectramax Plus 384) at 37°C.

### Folate assay

Folate was quantified using a microbiological assay as described (28, 68). *P. gingivalis* cells were centrifuged (12,000 g, 10 min, 20^0^C) then washed and resuspended in 0.1 M sodium acetate pH 4.8-1% ascorbic acid. Cells were disrupted in a FastPrep-24 5G (MP Biomedicals) and incubated at 100^0^C for 5 min to release folate from folate binding proteins. For deconjugation of polyglutamyl tails, 200 mg of human plasma (MilliporeSigma) was diluted in 1 ml of 0.1 M 2-mercaptoethanol-0.5% sodium ascorbate, and cleared from precipitates by centrifugation (10,000 g, 5 min). The clarified plasma solution was added to the *P. gingivalis* samples at 2.5% (vol/vol) and reacted for 4 h at 37^0^C. *L. rhamnosus* cells were adjusted to OD_600_ = 0.35, mixed with 200 volumes of 9 g/l NaCl, and 20 μl added to individual wells of a 96 well plate containing 8 μl of 0.8 M sodium ascorbate-50 mM potassium phosphate buffer, pH 6.1. *P. gingivalis* samples made up to 150 μl with distilled water, and 150 μl Folic Acid Medium (Himedia), were added to wells. Standards were 150 μl of folinic acid calcium salt hydrate (ThermoFisher). Reactions were incubated for 18 h at 37^0^C in 5% CO_2_. Bacterial clumps were dispersed by aspiration and OD_600_ measured in a Spectramax Plus 384.

### Human neutrophil isolation

Neutrophils were purified using plasma-Percoll gradients as previously described (69). The purity of the isolated cell fraction as determined by cytospins, Giemsa staining, and microscopic evaluation was ≥95%. Cell viability was ≥97% as determined by trypan blue exclusion. Neutrophils were routinely cultured in RPMI medium (Gibco) containing 10% heat-inactivated fetal bovine serum.

### Chemotaxis

Neutrophils (4 x 10^6^ cells/ml) were re-suspended in 1x HBSS media containing calcium chloride and magnesium chloride (Gibco). Neutrophils were reacted with *P. gingivalis* strains and pre-warmed in water bath at 37°C for 10 min. HBSS media containing 10 nM CXCL1 (Genscript) was added to the lower chamber of 24-well transwell plates containing a permeable insert, and 100 μl of pre-warmed neutrophils-bacterial cells mixture was added to the upper chamber. A separate well with only neutrophils in the upper chamber was used as a control. The plate was incubated at 37°C for 30 min in a 5% CO2 incubator. After 30 min, the transwell membranes were stained with a HEMA 3 stain set kit following the manufacturer’s instructions (ThermoFisher). Chemotaxis was assessed by light microscopic examination (magnification, x100) of the underside of the membrane. The average number of cells from a total of 10 fields was determined, and data were normalized by the area of the membrane circle and field of view.

### Opsonization

Bacteria (2 x 10^9^) were resuspended in 1 ml Hanks balanced salt solution (HBSS) (without Ca^2+^/Mg^2+^) and opsonized with 10% human serum, pooled from 10 healthy donors, for 1 h at 37°C with gentle rocking. Cells were stained with anti-C3b/iC3b conjugated with fluorescein isothiocyanate (FITC) (1:200), anti-IgG conjugated with phycoerythrin (PE) (1:100), or anti-IgM conjugated with allophycocyanin (APC) (1:50) (all from BioLegend) in flow buffer (PBS containing 2mM EDTA and 1% BSA) for 1 h at 4^0^C in the dark with rocking. After washing once with PBS by centrifugation at 5,000 g, 5min at 4^0^C, and fixing in 2% paraformaldehyde, data were acquired using a BD fluorescence-activated cell sorter and analyzed using FlowJo software.

### Phagocytosis

Neutrophils (4 × 10^6^ cells/ml) were reacted with serum-opsonized, CFSE-labeled *P. gingivalis* at MOI 10 in a shaking water bath at 37°C for 30 min. Cells were pelleted and rinsed at 600 g for 5 min. The neutrophil plasma membrane was stained for 10 min with Alexa Fluor 647 conjugated wheat germ agglutinin (Invitrogen) and fixed with 2% paraformaldehyde. Cells were analyzed using an Amnis ImageStream imaging cytometer (MilliporeSigma). Data were analyzed using IDEAS image data exploration and analysis software v.6.0 (Amnis), and quantitation of fluorescence was with ImageJ.

### Neutrophil killing assay

Opsonized bacteria were incubated (MOI 10) with neutrophils (4 × 10^6^ cells/ml) at 37°C for 30 min in shaking water bath. Neutrophils were washed twice in PBS by centrifugation at 100 g, 5 min, and internalized bacteria were released by hypotonic lysis in dH_2_O for 20 min. Serial dilutions of the lysates were plated on TSB blood agar and cultured anaerobically for CFU enumeration.

### Chemiluminescent detection of ROS

Neutrophils were suspended in PBS containing 90 mM calcium chloride, 50 mM magnesium chloride, 750 mM glucose, 125 μM luminol, and 100 U/mL horseradish peroxidase in 96-well white plates (Costar). Bacterial cells at MOI 10 were added, and total ROS production (measured in relative light units [RLU]) was recorded at 37°C continuously for 1 h in a Spectramax L luminometer. Intracellular ROS was detected by adding superoxide dismutase (SOD) (5 μL, 3 U/μL). As a positive control, neutrophils were stimulated with 175 nM PMA.

### MPO colocalization

Neutrophils (2 × 10^6^) were attached to serum treated coverslips in 24-well plates and reacted with CFSE-labeled bacteria (MOI 10). Phagocytosis was synchronized by centrifugation at 600 g for 4 min and proceeded for 1 h at 37°C, following which cells were fixed with 4% PFA and blocked with 1% BSA prepared in PBS without Ca^2+^ or Mg^2+^ for 1 h at RT. Cells were reacted with anti-A-LPS (1:1000) for 1 h at RT, washed with PBS and treated with Alexa fluor 555 conjugated goat anti-mouse IgG (1:2000) to detect extracellular bacteria. After 1 h incubation at RT, cells were washed and fixed using 4 % PFA for 8 min at room temperature. Cells were blocked and permeabilized with 1% BSA and 0.01% saponin in PBS for 1 h at RT. Cells were treated with rabbit anti-MPO (1:200) primary antibody overnight at 4°C, washed with the blocking solution before secondary antibody, Alexa fluor 647 goat anti-rabbit (1:2000), treatment for 2 h at RT. Following staining with DAPI, cells were analyzed by confocal microscopy. Images were processed using IMARIS software (Oxford Instruments).

### Neutrophil degranulation

Neutrophils (0.2 × 10^6^ per well) were suspended in RPMI in 96-well plates and stimulated with bacteria MOI 10 at 37^0^C for 4 h. Separately, neutrophils were stimulated with latrunculin (1 μM) for 30 min, followed by fMLF (N-formylmethionyl-leucyl-phenylalanine) (1 μM) for 10 min, to induce maximal degranulation. Elastase and MMP-9 release was measured in cell-free supernatants by ELISA (elastase/ELA2 DuoSet ELISA, or MMP9 Duoset ELISA, both R&D) and lactoferrin measured using the human ELISA kit (Abcam), following the manufacturers’ instructions.

### Neutrophil NET formation

Neutrophils (1 × 10^6^ cells/ml) were resuspended in prewarmed NET assay buffer (Abcam), distributed in 24-well plates, and reacted with bacteria at MOI 10, or with 20 nM phorbol myristate acetate (PMA) as a positive control, for 4 h at 37^0^C. Soluble, non-NET-associated elastase was removed by washing, and NETs were treated with S7 nuclease (15 U/mL) for 30 min at 37°C. After inactivation of nuclease with 500 nM EDTA and centrifugation to remove cell debris, neutrophil elastase substrate (1:1) was added, and the mixture was incubated for 3 h at 37°C. Absorbance was recorded at 405 nm.

### Ethics

Blood was drawn from healthy donors in accordance with the guidelines approved by the Institutional Review Board of the University of Louisville. Informed consent was obtained from each donor.

The University of Louisville Institutional Animal Care and Use Committee approved all animal procedures in this study, and all procedures were performed in accordance with the recommendations in the NIH *Guide for Care and Use of Laboratory Animals*. Mice were under the care of full-time staff, given access to food and water ad libitum and maintained on a 12-h light:dark cycle, with a temperature of 22-25^0^C and a relative humidity of 40–60%.

## Author contributions

RJL, KSS, JD, SMU, JB, DPM, JP conceived the study, analyzed data, wrote/edited the manuscript. SDP, SJ, JDP, LY, J-ZJ, HL, HK, DWW, ZRF, AV, IS, MKS performed experiments. MLM analyzed proteomics data.

## Supplementary Material

**Fig. S1.**
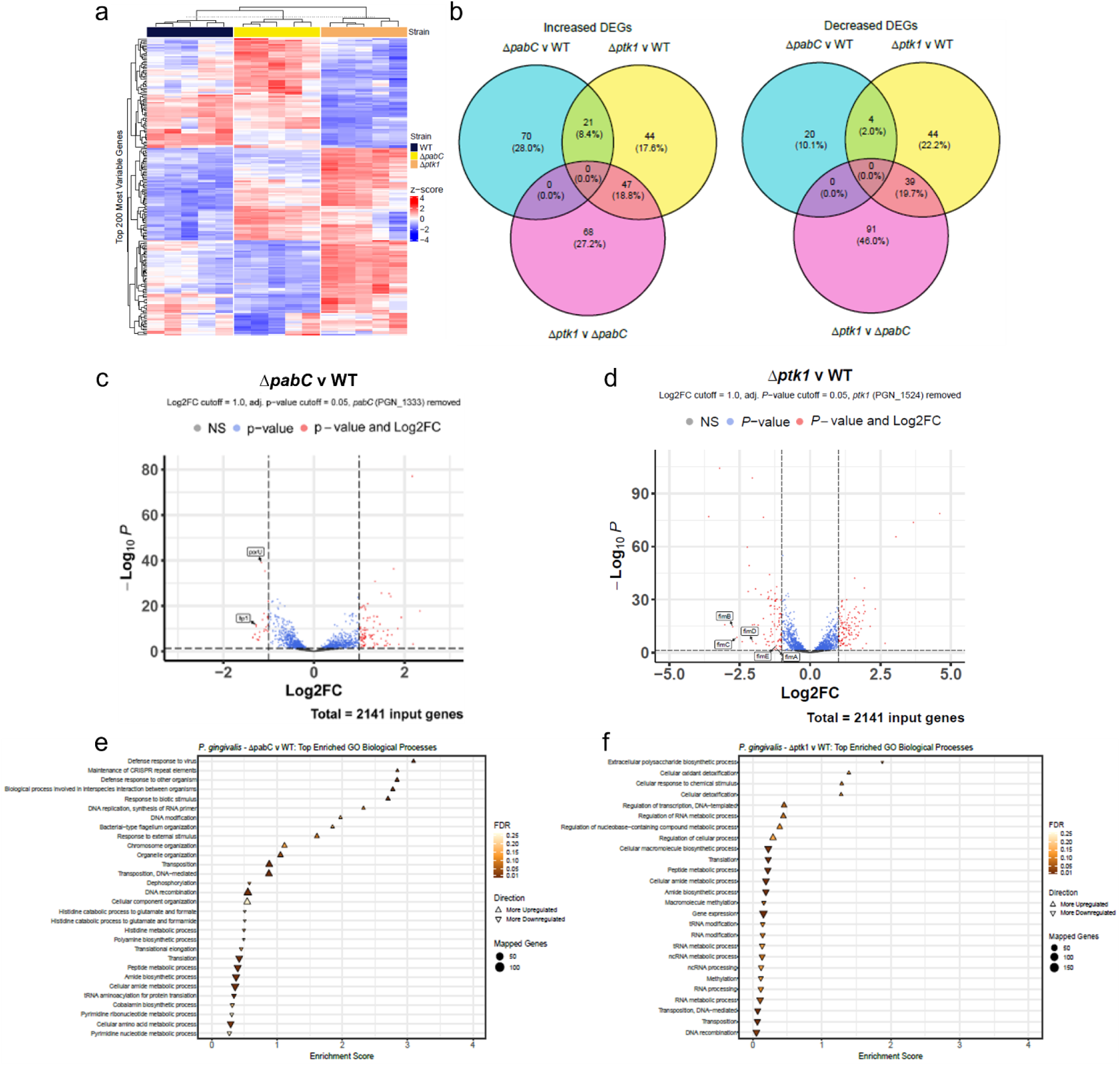
RNA-Seq analysis of differential mRNA expression in Δ*pabC* or Δ*ptk1*. a) Heat map showing z scores for top 200 differentially regulated genes. b) Venn diagrams of numbers of genes showing differential regulation. c) Volcano plot of differentially expressed genes in Δ*pabC* compared to WT. d) Volcano plot of differentially expressed genes in Δ*ptk1* compared to WT. e, f) GO Biological processes enrichment analysis of genes differentially regulated in e) Δ*pabC* or f) Δ*ptk1* as generated via STRING Database (V 12.0) using the “Proteins with Values/Ranks” function with an FDR of 25% and a confidence score of 0.4 for network generation.

**Fig S2.**
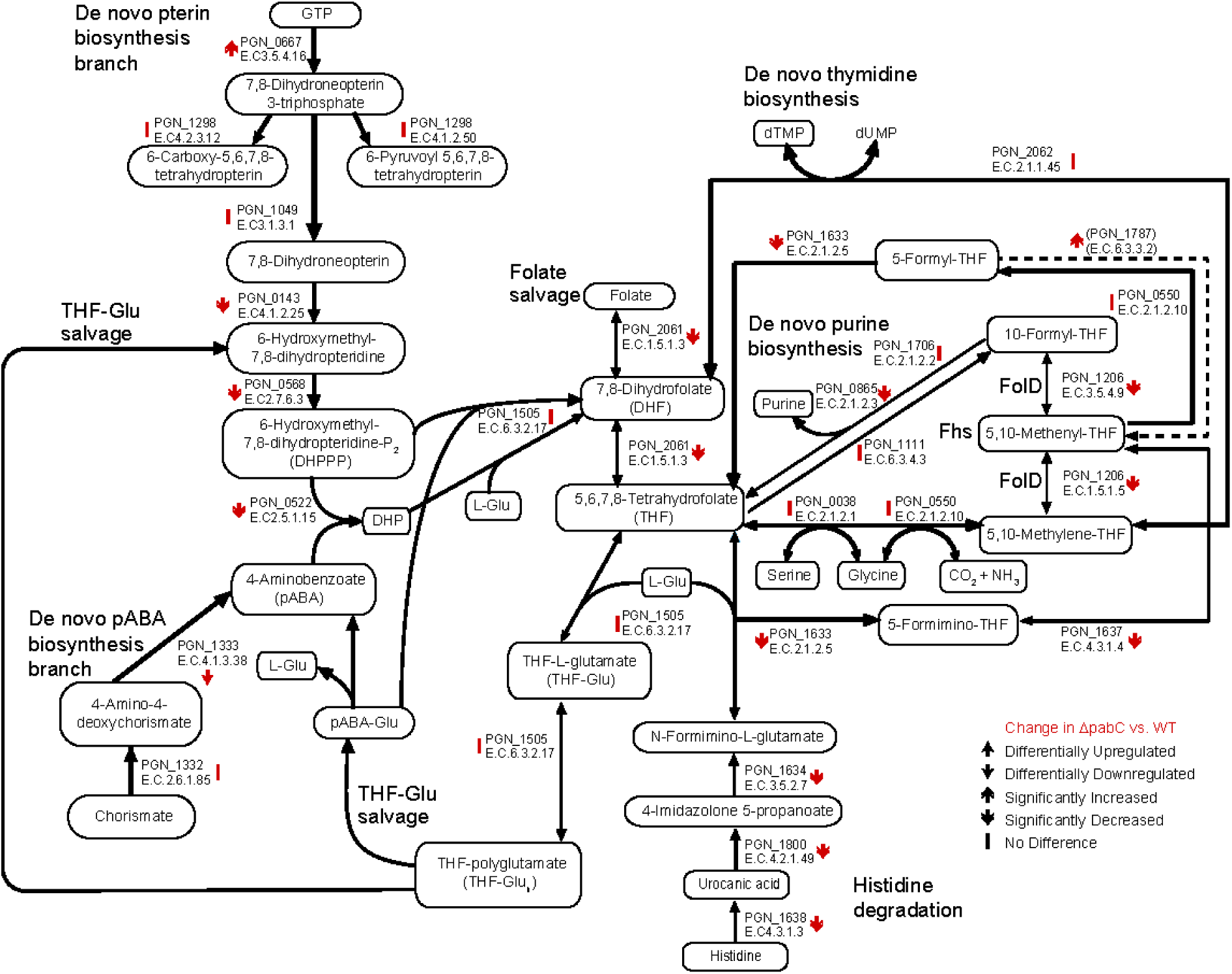
Representation of the OCM pathway showing genes differentially expressed in Δ*pabC* by RNA-Seq.

**Fig. S3.**
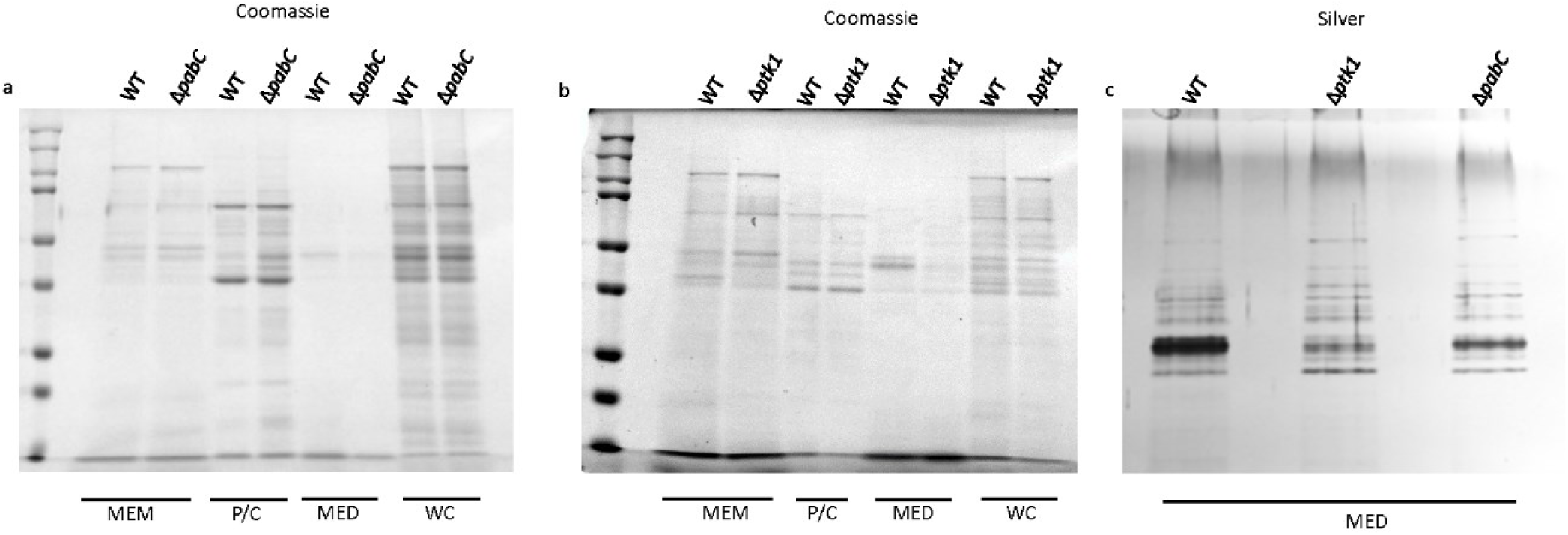
a, b) Coomassie blue stain of representative SDS-PAGE of *P. gingivalis* cell fractions analyzed in Fig 2. c) Silver stain of representative SDS-PAGE of *P. gingivalis* culture supernatants. The intense band is the catalytic domain from the gingipains.

**Fig S4.**
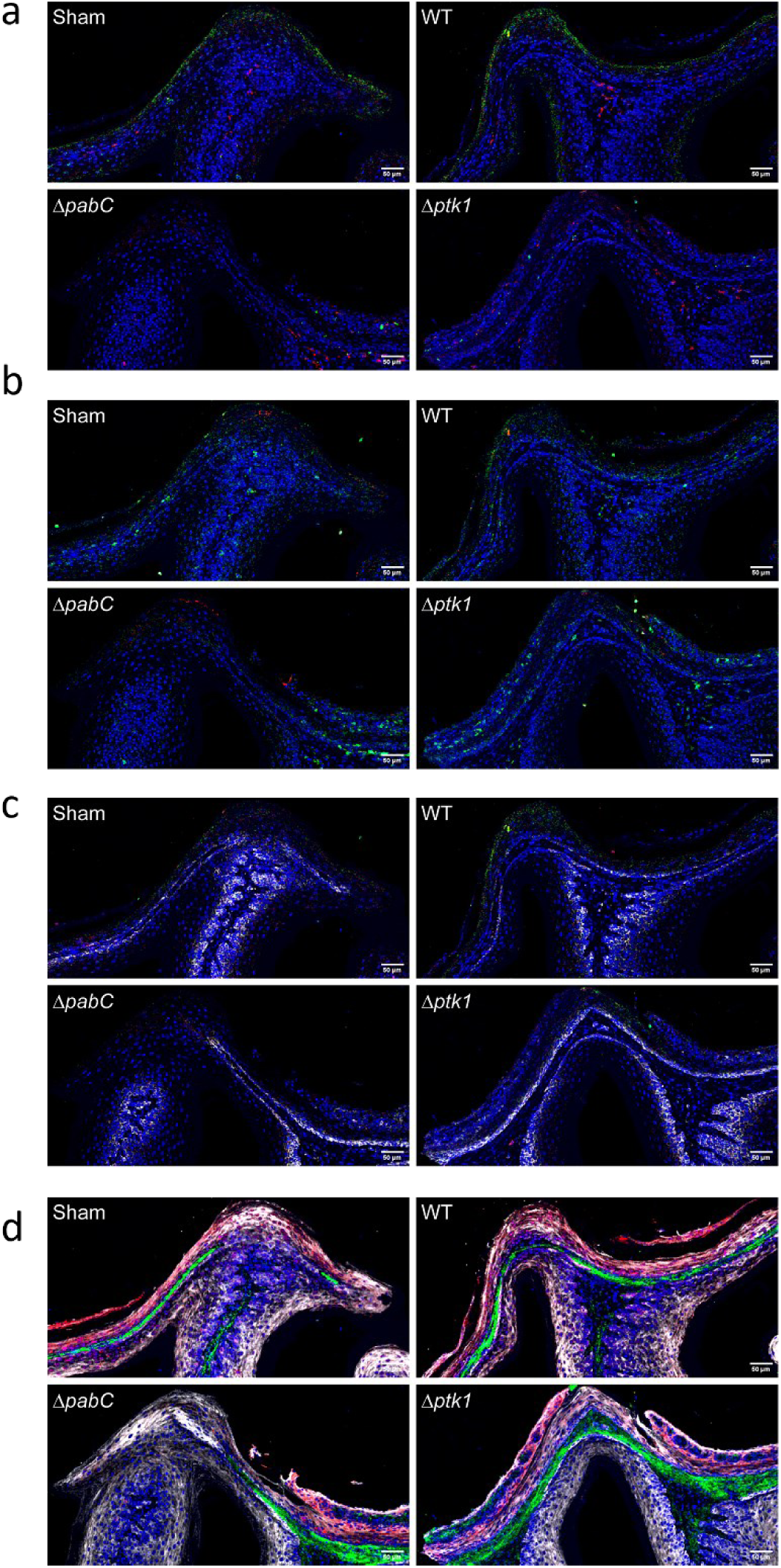
Representative mass cytometry images of gingival tissue from mice infected with WT, Δ*pabC*, Δ*ptk1*, or PBS (Sham) and stained DAPI and with antibodies recognizing: a) F480 (red) and MHCII (green); b) CD45 (green) and B220 (red); c) CD4 (red), CD8 (green), and CD44 (white); and d) collagen (green), PanCK (white), and EpCam (red). Bar = 50 μm.

**Fig. S5.**
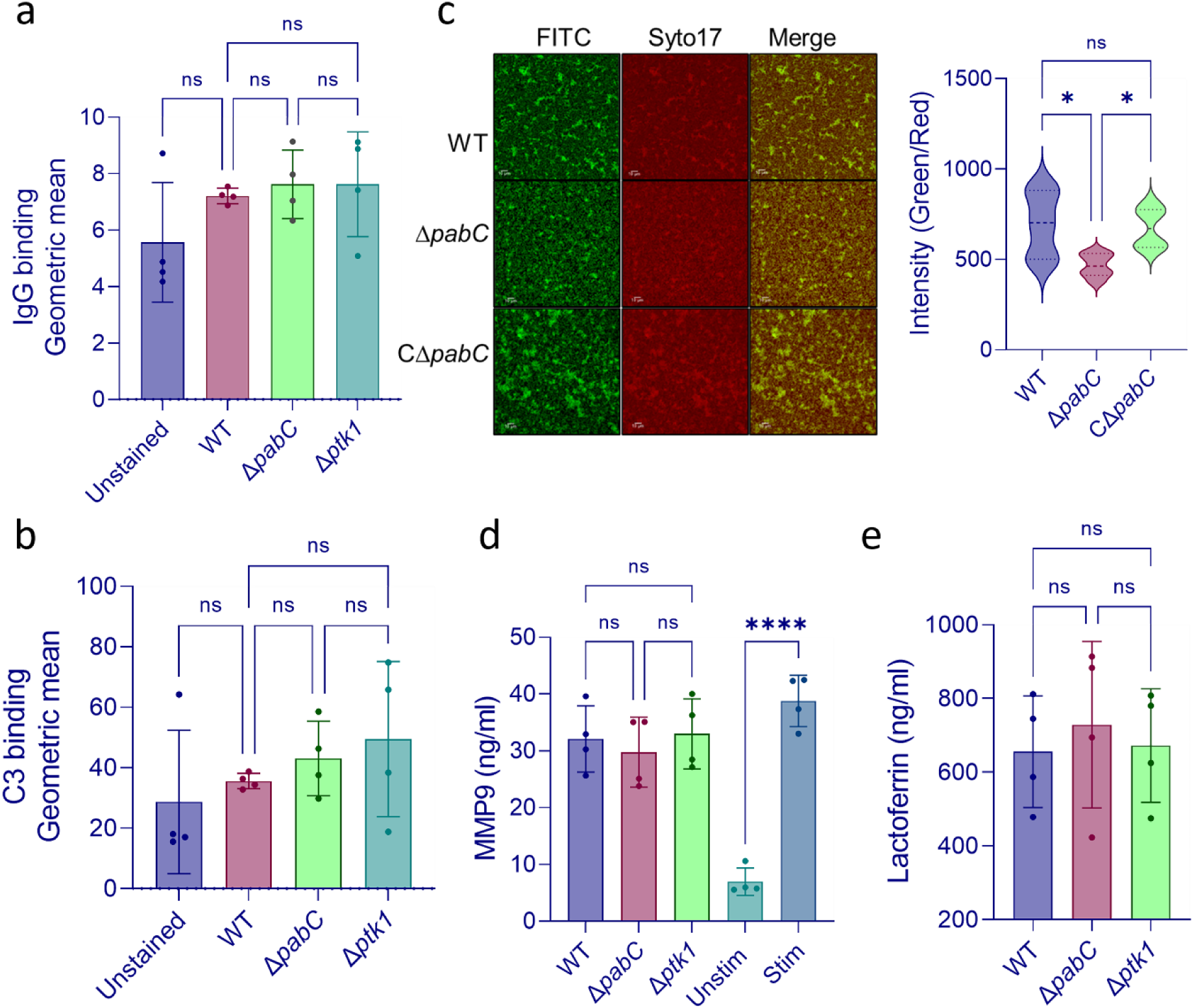
Properties of Δ*pabC* and Δ*ptk1* mutants. a-b) Opsonization of *P. gingivalis* strains indicated in 10% human serum. Bacteria were probed with conjugated with phycoerythrin (PE) conjugated IgG (a) or FITC-conjugated anti anti-C3b/iC3b (b) and analyzed by flow cytometry. Data are geometric means ± SD of fluorescence. c) EPS production by Δ*pabC.* Confocal microscopy of Syto 17 stained *P. gingivalis* probed with FITC-labeled Concanavalin A and WGA, and fluorescent intensities plotted. Data are expressed as fluorescent intensity ratios of lectins to bacterial cells. d) MMP9 and e) lactoferrin production from neutrophils challenged with serum-opsonized *P. gingivalis* strains MOI 10 for 4h. Unstim is unstimulated negative control and Stim is positive control with latrunculin and fMLF. Data are means +/- SD. \**P*<.05, ****P<.001 by ANOVA with Tukey’s correction for multiple comparisons. ns is not significant (*P*>.05). Data are representative of at least three independent experiments.

**Table S1.**
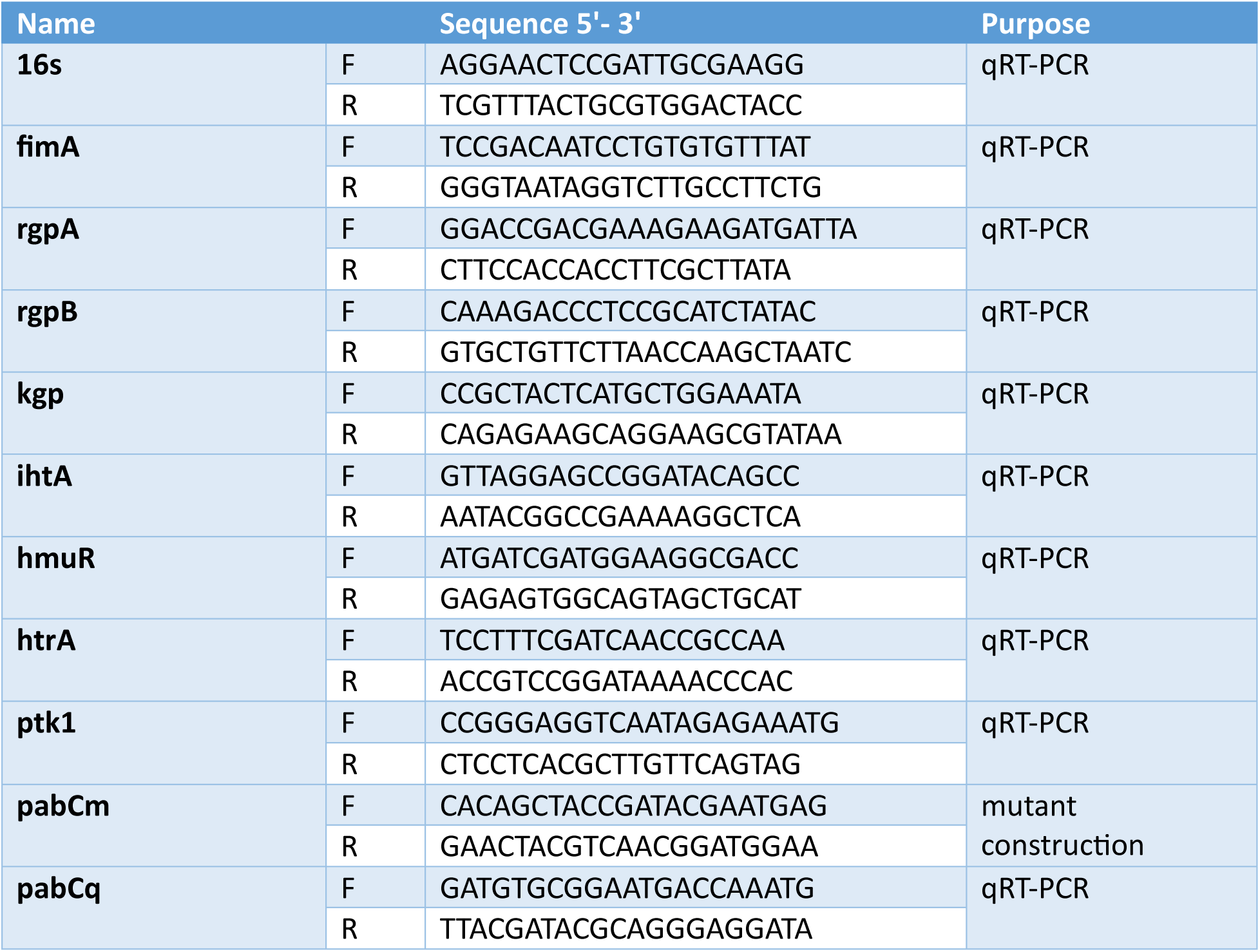
Primers used in this study.

**Table S2.** Antibodies used in IMC.

|  | Antibody | Cat# | Dilution |
| --- | --- | --- | --- |
| 1 | Anti-Collagen I (Polyclonal)-142Nd—15 µg; FFPE, Mouse; | 91H018142 | 1:400 |
| 2 | Anti-Hu/Ms CD11b (EPR1344)-146Nd—15 µg | 91H007146 | 1:200 |
| 3 | Anti-Ki-67 (B56)-150Nd—15 µg; FFPE, Human, Mouse; | 91H017150 | 1:200 |
| 4 | Anti-CD45 (D3F8Q)-156Gd—15 µg | 91H029156 | 1:200 |
| 5 | Anti-Ly6G (1A8)-160Gd—15 µg; FFPE, Mouse | 91H037160 | 1:200 |
| 6 | Anti-Human/Mouse CD44 (IM7)-162Dy—100 Tests | 3162030B | 1:400 |
| 7 | Anti-Mouse CX3CR1 (SA011F11)-164Dy—100 Tests | 3164023B | 1:200 |
| 8 | Anti-CD4 (BLR167J)-165Ho—15 µg; FFPE, Mouse | 91H031165 | 1:200 |
| 9 | Anti-F4/80 (D2S9R)-166Er—15 µg, FFPE, Mouse | 91H030166 | 1:200 |
| 10 | Anti-CD8 (EPR21769)-167Er—15 µg; FFPE, Mouse | 91H023167 | 1:200 |
| 11 | Anti-MHC-II (M5/114.15.2)-169Tm—15 µg; FFPE, Mouse | 91H038169 | 1:200 |
| 12 | Anti-CD31/PECAM (EPR17259)-171Yb—15 µg, FFPE, | 91H027171 | 1:300 |
| 13 | Anti-Pan-CytoKeratin (AE-1/AE-3)-174Yb—15 µg; FFPE, Mouse; | 91H006174 | 1:300 |

**Table S3.** Recombinant Protein Expression.

| Protein | Vector | Purification | Concentration (mg/mL) | Confirmed by LC-MS | Purity Method | Purity | Storage Buffer |
| --- | --- | --- | --- | --- | --- | --- | --- |
| **PGN_1049_His** | pET30a | Supernatant of cell lysate, Ni column | 0.66 | Yes | SDS-PAGE under reducing conditions | >= 55% | 50 mM Tris-HCL, 150 mM NaCl, 10% Glycerol, 1 mM TCEP, pH 8.0 |
| **PGN_0550_His** | pET30a | Supernatant of cell lysate, Ni column | 8.34 | Yes | SDS-PAGE under reducing conditions | >= 90% | 50 mM Tris-HCL, 150 mM NaCl, 10% Glycerol, 1 mM TCEP, pH 8.0 |
| **PGN_0038_His** | pET30a | Supernatant of cell lysate, Ni column | 7.18 | Yes | SDS-PAGE under reducing conditions | >= 90% | 50 mM Tris-HCL, 150 mM NaCl, 10% Glycerol, pH 8.0 |
| **PGN_0522_His** | pET30a | Supernatant of cell lysate, Ni column | 3.20 | Yes | SDS-PAGE under reducing conditions | >= 90% | 50 mM Tris-HCL, 150 mM NaCl, 10% Glycerol, pH 8.0 |

## References

1. Berberolli S, Wu M, Goycoolea FM. 2024. The Rosetta Stone of interactions of mucosa and associated bacteria in the gastrointestinal tract. Curr Opin Gastroenterol. 40:1–6.

2. Chen Y, Quirk NF, Tan S. 2023. Shining a light on bacterial environmental cue integration and its relation to metabolism. Mol Microbiol. 120:71–74.

3. Pokorzynski ND, Groisman EA. 2023. How bacterial pathogens coordinate appetite with virulence. Microbiol Mol Biol Rev. 87:e0019822.

4. Stocke KS, Lamont RJ. 2023. One-carbon metabolism and microbial pathogenicity. Mol Oral Microbiol. 39:156–164.

5. Shetty S, Varshney U. 2021. Regulation of translation by one-carbon metabolism in bacteria and eukaryotic organelles. J Biol Chem. 296:100088.

6. Chimalapati S, Cohen J, Camberlein E, Durmort C, Baxendale H, de Vogel C, van Belkum A, Brown JS. 2011. Infection with conditionally virulent *Streptococcus pneumoniae* Δ*pab* strains induces antibody to conserved protein antigens but does not protect against systemic infection with heterologous strains. Infect Immun. 79:4965–4976.

7. Brown JS, Aufauvre-Brown A, Brown J, Jennings JM, Arst H, Jr., Holden DW. 2000. Signature-tagged and directed mutagenesis identify PABA synthetase as essential for *Aspergillus fumigatus* pathogenicity. Mol Microbiol. 36:1371–1380.

8. Dial CN, Speare L, Sharpe GC, Gifford SM, Septer AN, Visick KL. 2021. Para-Aminobenzoic Acid, Calcium, and c-di-GMP Induce Formation of Cohesive, Syp-Polysaccharide-Dependent Biofilms in *Vibrio fischeri*. mBio. 12:e0203421.

9. Kuboniwa M, Houser JR, Hendrickson EL, Wang Q, Alghamdi SA, Sakanaka A, Miller DP, Hutcherson JA, Wang T, Beck DAC, Whiteley M, Amano A, Wang H, Marcotte EM, Hackett M, Lamont RJ. 2017. Metabolic crosstalk regulates *Porphyromonas gingivalis* colonization and virulence during oral polymicrobial infection. Nat Microbiol. 2:1493–1499.

10. Hajishengallis G, Liang S, Payne MA, Hashim A, Jotwani R, Eskan MA, McIntosh ML, Alsam A, Kirkwood KL, Lambris JD, Darveau RP, Curtis MA. 2011. Low-abundance biofilm species orchestrates inflammatory periodontal disease through the commensal microbiota and complement. Cell Host Microbe. 10:497–506.

11. Green JM, Nichols BP. 1991. p-Aminobenzoate biosynthesis in *Escherichia coli*. Purification of aminodeoxychorismate lyase and cloning of pabC. J Biol Chem. 266:12971–12975.

12. Zhou M, Ji M. 2005. Molecular docking and 3D-QSAR on 2-(oxalylamino) benzoic acid and its analogues as protein tyrosine phosphatase 1B inhibitors. Bioorg Med Chem Lett. 15:5521–5525.

13. Jung YJ, Miller DP, Perpich JD, Fitzsimonds ZR, Shen D, Ohshima J, Lamont RJ. 2019. *Porphyromonas gingivalis* Tyrosine Phosphatase Php1 Promotes Community Development and Pathogenicity. mBio. 10:e02004–02019.

14. Wright CJ, Xue P, Hirano T, Liu C, Whitmore SE, Hackett M, Lamont RJ. 2014. Characterization of a bacterial tyrosine kinase in *Porphyromonas gingivalis* involved in polymicrobial synergy. Microbiologyopen. 3:383–394.

15. Shen D, Perpich JD, Stocke KS, Yakoumatos L, Fitzsimonds ZR, Liu C, Miller DP, Lamont RJ. 2020. Role of the RprY response regulator in *P. gingivalis* community development and virulence. Mol Oral Microbiol. 35:231–239.

16. Tjaden B. 2020. A computational system for identifying operons based on RNA-seq data. Methods. 176:62–70.

17. Miller DP, Hutcherson JA, Wang Y, Nowakowska ZM, Potempa J, Yoder-Himes DR, Scott DA, Whiteley M, Lamont RJ. 2017. Genes contributing to *Porphyromonas gingivalis* fitness in abscess and epithelial cell colonization environments. Front Cell Infect Microbiol. 7:378.

18. Lamont RJ, Miller DP. 2022. Tyrosine Kinases and Phosphatases: Enablers of the *Porphyromonas gingivalis* Lifestyle. Front Oral Health. 3:835586.

19. Irfan M, Solbiati J, Duran-Pinedo A, Rocha FG, Gibson FC, 3rd, Frias-Lopez J. 2024. A *Porphyromonas gingivalis* hypothetical protein controlled by the type I-C CRISPR-Cas system is a novel adhesin important in virulence. mSystems. 9:e0123123.

20. Nowakowska Z, Madej M, Grad S, Wang T, Hackett M, Miller DP, Lamont RJ, Potempa J. 2021. Phosphorylation of major *Porphyromonas gingivalis* virulence factors is crucial for their processing and secretion. Mol Oral Microbiol. 36:316–326.

21. Lasica AM, Goulas T, Mizgalska D, Zhou X, de Diego I, Ksiazek M, Madej M, Guo Y, Guevara T, Nowak M, Potempa B, Goel A, Sztukowska M, Prabhakar AT, Bzowska M, Widziolek M, Thogersen IB, Enghild JJ, Simonian M, Kulczyk AW, Nguyen KA, Potempa J, Gomis-Ruth FX. 2016. Structural and functional probing of PorZ, an essential bacterial surface component of the type-IX secretion system of human oral-microbiomic *Porphyromonas gingivalis*. Sci Rep. 6:37708.

22. Gorasia DG, Veith PD, Chen D, Seers CA, Mitchell HA, Chen YY, Glew MD, Dashper SG, Reynolds EC. 2015. *Porphyromonas gingivalis* Type IX Secretion Substrates Are Cleaved and Modified by a Sortase-Like Mechanism. PLoS Pathog. 11:e1005152.

23. Uriarte SM, Hajishengallis G. 2023. Neutrophils in the periodontium: Interactions with pathogens and roles in tissue homeostasis and inflammation. Immunol Rev. 314:93–110.

24. Sochalska M, Potempa J. 2017. Manipulation of Neutrophils by *Porphyromonas gingivalis* in the Development of Periodontitis. Front Cell Infect Microbiol. 7:197.

25. Shoji M, Sato K, Yukitake H, Naito M, Nakayama K. 2014. Involvement of the Wbp pathway in the biosynthesis of *Porphyromonas gingivalis* lipopolysaccharide with anionic polysaccharide. Sci Rep. 4:5056.

26. Chen T, Siddiqui H, Olsen I. 2017. In silico Comparison of 19 *Porphyromonas gingivalis* Strains in Genomics, Phylogenetics, Phylogenomics and Functional Genomics. Front Cell Infect Microbiol. 7:28.

27. Nadkarni MA, Chhour KL, Browne G, Jacques NA, Hunter N. 2009. Lysine gingipain (*kgp*) biovars of *Porphyromonas gingivalis* exhibit differential distribution on oral mucosal sites. J Clin Microbiol. 47:3350–3352.

28. Sybesma W, Starrenburg M, Kleerebezem M, Mierau I, de Vos WM, Hugenholtz J. 2003. Increased production of folate by metabolic engineering of *Lactococcus lactis*. Appl Environ Microbiol. 69:3069–3076.

29. Sybesma W, Van Den Born E, Starrenburg M, Mierau I, Kleerebezem M, De Vos WM, Hugenholtz J. 2003. Controlled modulation of folate polyglutamyl tail length by metabolic engineering of *Lactococcus lactis*. Appl Environ Microbiol. 69:7101–7107.

30. Nemoto TK, Ohara-Nemoto Y. 2016. Exopeptidases and gingipains in *Porphyromonas gingivalis* as prerequisites for its amino acid metabolism. Jpn Dent Sci Rev. 52:22–29.

31. Kalhan SC, Hanson RW. 2012. Resurgence of serine: an often neglected but indispensable amino Acid. J Biol Chem. 287:19786–19791.

32. Hajredini F, Alphonse S, Ghose R. 2023. BY-kinases: Protein tyrosine kinases like no other. J Biol Chem. 299:102737.

33. Robertson CD, Hazen TH, Kaper JB, Rasko DA, Hansen AM. 2018. Phosphotyrosine-Mediated Regulation of Enterohemorrhagic *Escherichia coli* Virulence. mBio. 9.

34. Grangeasse C. 2016. Rewiring the Pneumococcal Cell Cycle with Serine/Threonine- and Tyrosine-kinases. Trends Microbiol. 24:713–724.

35. Mijakovic I, Grangeasse C, Turgay K. 2016. Exploring the diversity of protein modifications: special bacterial phosphorylation systems. FEMS Microbiol Rev. 40:398–417.

36. Alphonse S, Djemil I, Piserchio A, Ghose R. 2022. Structural basis for the recognition of the bacterial tyrosine kinase Wzc by its cognate tyrosine phosphatase Wzb. Proc Natl Acad Sci U S A. 119:e2201800119.

37. Lasica AM, Ksiazek M, Madej M, Potempa J. 2017. The Type IX Secretion System (T9SS): Highlights and Recent Insights into Its Structure and Function. Front Cell Infect Microbiol. 7:215.

38. Veith PD, Glew MD, Gorasia DG, Cascales E, Reynolds EC. 2022. The Type IX Secretion System and Its Role in Bacterial Function and Pathogenesis. J Dent Res. 101:374–383.

39. Mizgalska D, Goulas T, Rodriguez-Banqueri A, Veillard F, Madej M, Malecka E, Szczesniak K, Ksiazek M, Widziolek M, Guevara T, Eckhard U, Sola M, Potempa J, Gomis-Ruth FX. 2021. Intermolecular latency regulates the essential C-terminal signal peptidase and sortase of the *Porphyromonas gingivalis* type-IX secretion system. Proc Natl Acad Sci U S A. 118.

40. Salaun C, Greaves J, Chamberlain LH. 2010. The intracellular dynamic of protein palmitoylation. J Cell Biol. 191:1229–1238.

41. Fraser JA, Worrall EG, Lin Y, Landre V, Pettersson S, Blackburn E, Walkinshaw M, Muller P, Vojtesek B, Ball K, Hupp TR. 2015. Phosphomimetic mutation of the N-terminal lid of MDM2 enhances the polyubiquitination of p53 through stimulation of E2-ubiquitin thioester hydrolysis. J Mol Biol. 427:1728–1747.

42. Bochner BR, Ames BN. 1982. ZTP (5-amino 4-imidazole carboxamide riboside 5’-triphosphate): a proposed alarmone for 10-formyl-tetrahydrofolate deficiency. Cell. 29:929–937.

43. Sybesma W, Starrenburg M, Tijsseling L, Hoefnagel MH, Hugenholtz J. 2003. Effects of cultivation conditions on folate production by lactic acid bacteria. Appl Environ Microbiol. 69:4542–4548.

44. Lionaki E, Ploumi C, Tavernarakis N. 2022. One-Carbon Metabolism: Pulling the Strings behind Aging and Neurodegeneration. Cells. 11.

45. Lamont RJ, Koo H, Hajishengallis G. 2018. The oral microbiota: dynamic communities and host interactions. Nat Rev Microbiol. 16:745–759.

46. Daep CA, Novak EA, Lamont RJ, Demuth DR. 2011. Structural dissection and in vivo effectiveness of a peptide inhibitor of *Porphyromonas gingivalis* adherence to *Streptococcus gordonii*. Infect Immun. 79:67–74.

47. Kin LX, Butler CA, Slakeski N, Hoffmann B, Dashper SG, Reynolds EC. 2020. Metabolic cooperativity between *Porphyromonas gingivalis* and *Treponema denticola*. J Oral Microbiol. 12:1808750.

48. Lamont RJ, Jenkinson HF. 1998. Life below the gum line: pathogenic mechanisms of *Porphyromonas gingivalis*. Microbiol Mol Biol Rev. 62:1244–1263.

49. Lee JY, Miller DP, Wu L, Casella CR, Hasegawa Y, Lamont RJ. 2018. Maturation of the Mfa1 Fimbriae in the Oral Pathogen *Porphyromonas gingivalis*. Front Cell Infect Microbiol. 8:137.

50. Simionato MR, Tucker CM, Kuboniwa M, Lamont G, Demuth DR, Tribble GD, Lamont RJ. 2006. *Porphyromonas gingivalis* genes involved in community development with *Streptococcus gordonii*. Infect Immun. 74:6419–6428.

51. Potempa J, Pike R, Travis J. 1995. The multiple forms of trypsin-like activity present in various strains of *Porphyromonas gingivalis* are due to the presence of either Arg-gingipain or Lys-gingipain. Infect Immun. 63:1176–1182.

52. Nguyen KA, Travis J, Potempa J. 2007. Does the importance of the C-terminal residues in the maturation of RgpB from *Porphyromonas gingivalis* reveal a novel mechanism for protein export in a subgroup of Gram-Negative bacteria? J Bacteriol. 189:833–843.

53. Sztukowska M, Sroka A, Bugno M, Banbula A, Takahashi Y, Pike RN, Genco CA, Travis J, Potempa J. 2004. The C-terminal domains of the gingipain K polyprotein are necessary for assembly of the active enzyme and expression of associated activities. Mol Microbiol. 54:1393–1408.

54. Regnier P, Marques C, Saadoun D. 2023. PICAFlow: a complete R workflow dedicated to flow/mass cytometry data, from pre-processing to deep and comprehensive analysis. Bioinform Adv. 3:vbad177.

55. Levine JH, Simonds EF, Bendall SC, Davis KL, Amir el AD, Tadmor MD, Litvin O, Fienberg HG, Jager A, Zunder ER, Finck R, Gedman AL, Radtke I, Downing JR, Pe’er D, Nolan GP. 2015. Data-Driven Phenotypic Dissection of AML Reveals Progenitor-like Cells that Correlate with Prognosis. Cell. 162:184–197.

56. Chen S. 2023. Ultrafast one-pass FASTQ data preprocessing, quality control, and deduplication using fastp. Imeta. 2:e107.

57. Kim D, Paggi JM, Park C, Bennett C, Salzberg SL. 2019. Graph-based genome alignment and genotyping with HISAT2 and HISAT-genotype. Nat Biotechnol. 37:907–915.

58. Liao Y, Smyth GK, Shi W. 2014. featureCounts: an efficient general purpose program for assigning sequence reads to genomic features. Bioinformatics. 30:923–930.

59. Love MI, Huber W, Anders S. 2014. Moderated estimation of fold change and dispersion for RNA-seq data with DESeq2. Genome Biol. 15:550.

60. Benjamini Y, Drai D, Elmer G, Kafkafi N, Golani I. 2001. Controlling the false discovery rate in behavior genetics research. Behav Brain Res. 125:279–284.

61. Zhu A, Ibrahim JG, Love MI. 2019. Heavy-tailed prior distributions for sequence count data: removing the noise and preserving large differences. Bioinformatics. 35:2084–2092.

62. Szklarczyk D, Kirsch R, Koutrouli M, Nastou K, Mehryary F, Hachilif R, Gable AL, Fang T, Doncheva NT, Pyysalo S, Bork P, Jensen LJ, von Mering C. 2023. The STRING database in 2023: protein-protein association networks and functional enrichment analyses for any sequenced genome of interest. Nucleic Acids Res. 51:D638–D646.

63. Wickham H. 2016. ggplot2: elegant graphics for data analysis, 2nd ed.. Springer-Verlag, New York, NY.

64. Pedersen T. 2021, posting date. Ggforce: accelerating ‘ggplot2’. R package version 0.3.3. [Online.]

65. Gu Z. 2022. Complex heatmap visualization. Imeta. 1:e43.

66. Caster DJ, Korte EA, Merchant ML, Klein JB, Wilkey DW, Rovin BH, Birmingham DJ, Harley JB, Cobb BL, Namjou B, McLeish KR, Powell DW. 2015. Autoantibodies targeting glomerular annexin A2 identify patients with proliferative lupus nephritis. Proteomics Clin Appl. 9:1012–1020.

67. Grangeasse C, Doublet P, Cozzone AJ. 2002. Tyrosine phosphorylation of protein kinase Wzc from *Escherichia coli* K12 occurs through a two-step process. J Biol Chem. 277:7127–7135.

68. Horne DW, Patterson D. 1988. *Lactobacillus casei* microbiological assay of folic acid derivatives in 96-well microtiter plates. Clin Chem. 34:2357–2359.

69. Uriarte SM, Rane MJ, Luerman GC, Barati MT, Ward RA, Nauseef WM, McLeish KR. 2011. Granule exocytosis contributes to priming and activation of the human neutrophil respiratory burst. J Immunol. 187:391–400.

